# Dorsal raphe nucleus to anterior cingulate cortex 5-HTergic neural circuit modulates consolation and sociability

**DOI:** 10.1101/2020.09.21.307280

**Authors:** Lai-Fu Li, Li-Zi Zhang, Zhi-Xiong He, Yu-Ting Zhang, Huan Ma, Yu-Feng Xun, Wei Yuan, Wen-Juan Hou, Yi-Tong Li, Zi-Jian Lv, Rui Jia, Fa-Dao Tai

**Author notes:** Corresponding authors: Fa-Dao Tai, PhD, Professor, College of Life Sciences, Shaanxi Normal University, Xi’an, 710062, China. The first two authors contributed equally to this article.

## Abstract

Consolation is a common response to the distress of others in humans and some social animals, but the neural mechanisms underlying this behavior are not well characterized. By using socially monogamous mandarin voles, we found that optogenetic or chemogenetic inhibition of 5-HTergic neurons in the dorsal raphe nucleus (DR) or optogenetic inhibition of 5-HT terminals in the anterior cingulate cortex (ACC) significantly decreased allogrooming time in the consolation test and reduced sociability in the three-chamber test. The release of 5-HT within the ACC and the activity of DR neurons were significantly increased during allogrooming, sniffing and social approaching. Finally, we found that the activation of 5-HT1A receptors in the ACC was sufficient to reverse consolation and sociability deficits induced by the chemogenetic inhibition of 5-HTergic neurons in the DR. Our study provided first direct evidence that DR-ACC 5-HTergic neural circuit is implicated in consolation-like behaviors and sociability.

## INTRODUCTION

Consolation behavior, which is referred to as an increase in affiliative contact toward a distressed individual by an uninvolved bystander, is an important component of the social capabilities of humans (de Waal and Preston, 2017; Field et al., 2009). Impaired consolation has been frequently observed in many psychiatric diseases, such as depression, autism, and schizophrenia (Young et al., 2015). According to de Waal’s multilevel conceptualization of empathy (de Waal, 2008; de Waal and Preston, 2017), consolation represent an intermediate level of empathy (the primary level of “emotional contagion”, the more complex level of “consolation”, and the most elaborate level of “perspective taking and targeted helping”), which has long been assumed to exist in species possessing complex cognitive functions, such as humans, apes, dolphins and elephants (Perez-Manrique and Gomila, 2018); however, recent studies have indicated that it also exists in some socially lived rodents, such as prairie voles (Burkett et al., 2016), mandarin voles (Li et al., 2019) and rats (Knapska et al., 2010).

Currently, studies of the neural mechanisms underlying consolation and other forms of empathy have primarily focused on the oxytocin systems (Burkett et al., 2016; Li et al., 2019). However, as a complex social behavior, consolation may require the coordinated actions of numerous neuromodulators and neurotransmitters. Serotonin (5-HT) is an evolutionarily ancient neurotransmitter that has long been implicated in a variety of emotional disorders (Faye et al., 2020; Garcia-Garcia et al., 2018; Meneses and Liy-Salmeron, 2012). According to recent studies, 5-HT transmission is also involved in a series of social behaviors such as social interaction (Walsh et al., 2018), social reward and aggression (Dolen et al., 2013). Regarding empathy, a recent study revealed an association between salivary 5-HT levels and the empathic abilities of people (Matsunaga et al., 2017); a polymorphism in the promoter region of the serotonin transporter gene has been linked to individual differences in empathy (Gyurak et al., 2013); and MDMA (±3,4-methylenedioxymethamphetamine, better known as the recreational drug ‘‘ecstasy’’), which is well known to stimulate a feeling of closeness and empathy in its users (Carlyle et al., 2019), had been confirmed to robustly increase the release of 5-HT in an activity-independent manner (Heifets and Malenka, 2016). In animal studies, Kim, et al. found that microinjection of 5-HT into the anterior cingulate cortex (ACC) impairs vicarious fear and alters the regularity of neural oscillations in mice (Kim et al., 2014). Our recent study indicated that 5-HT1A receptors within the ACC are involved in consolation deficits induced by chronic social defeat stress in mandarin voles (Li et al., 2020). However, to our knowledge, direct evidence for an association between 5-HT and consolation has yet to be obtained.

Dorsal raphe nucleus (DR) is a main source of 5-HT neurons and provide 70% of 5-HTergic projections in the forebrain (Fu et al., 2010; Luo et al., 2015). DR 5-HTergic neurons form dense, broad and bidirectional neural connections with a broad range of forebrain and limbic structures, including the ACC (Celada et al., 2013; Charnay and Leger, 2010), which is a central hub for various types of empathy. Therefore, direct modulation of the DR→ACC 5-HTergic circuit to investigate its function role in consolation-like behaviors is interesting and meaningful.

The released 5-HT binds to pre- and postsynaptic receptors. To date, at least 14 different 5-HT receptor subtypes have been identified in the brain (Artigas, 2013). Among which, 5HT1AR and 5HT2AR are the two main subtypes that are expressed at high levels in the prefrontal cortex (Carhart-Harris and Nutt, 2017; Santana and Artigas, 2017). The distribution, signaling pathways and functions of these two receptors are substantially different, and both receptors play critical roles in modulating cortical activity and neural oscillations (Celada et al., 2013). Previous studies indicated that 5HT2AR gene single nucleotide polymorphisms are associated with empathy-related social communication abilities (Gong et al., 2015), and a 5HT2AR agonist increases emotional empathic ability (Dolder et al., 2017). However, in animal studies by Kim, et al., blockade of serotonin receptors in the ACC did not affect observational fear responses in mice (Kim et al., 2014). Clearly, the specific functions of 5HT1AR and 5HT2AR in empathy-like behaviors still require further examination.

The mandarin vole (*Microtus mandarinus*) is a socially monogamous rodent that is widely distributed across China (He et al., 2019). As shown in our previous studies, this species is capable of displaying consolation-like behaviors upon exposure to a distressed partner (Li et al., 2020; Li et al., 2019). In the present study, we first investigated the function of DR→ACC 5-HTergic circuits in consolation-like behavior using optogenetic and chemogenetic approaches. To provide more direct evidence, we then monitored ACC 5-HT release and DR neuron activities during this behavior by using *in vivo* fiber photometry. Finally, we used chemogenetics plus pharmacological approaches to investigate which types of 5-HT receptors in the ACC are involved in consolation-like behaviors in mandarin voles. In order to investigate any potential sex differences during these processes, both male and female subjects were included in our study. As consolation is in general a pro-social behavior, some social behaviors were investigated during the tests.

## RESULTS

### Optogenetic inhibition of DR 5-HT neurons in the DR→ACC neural circuit impaired consolation and reduced sociability

We first determined the 5-HTergic projection relationship between the DR and ACC in mandarin voles. For this experiment, the retrograde tracer CTB was injected into the ACC, followed by immunofluorescence staining of the DR sections with TPH2, a marker of 5-HTergic neurons. A substantial number of TPH2+ neurons colocalized with CTB, indicating the presence of DR-ACC 5-HTergic projections (Figure 1—figure supplement 1). We then used a novel dual-virus optogenetics approach to explore the function of the DR-ACC 5-HTergic circuit in consolation-like behaviors and sociability, where double-floxed AAV-DIO-ChR2-mCherry (DIO-ChR2) or AAV-DIO-eNpHR3.0-mCherry (DIO-eNpHR3.0) was injected into the DR and retro-AAVs containing the TPH2 promoter and Cre element (rAAV(Retro)-TPH2-Cre) were injected into the ACC (Figure 1A). This virus strategy ensures that opsins (excitatory CHR2 or inhibitory eNpHR3.0) are mainly expressed within the DR-ACC 5-HTergic circuit. Immunohistochemical staining showed that more than 75% of mCherry-labeled neurons expressed TPH2 in both male and female voles, and more than 80% of TPH2+ cells coexpressed mCherry (Figure 1—figure supplement 2). In the electrophysiological study, we found that the DR neurons reliably responded to pulses of 473 (activation)/593 (inhibition) nm light stimuli (Figure 1D-E). These results indicate the viability of this virus strategy.

**Figure 1.**
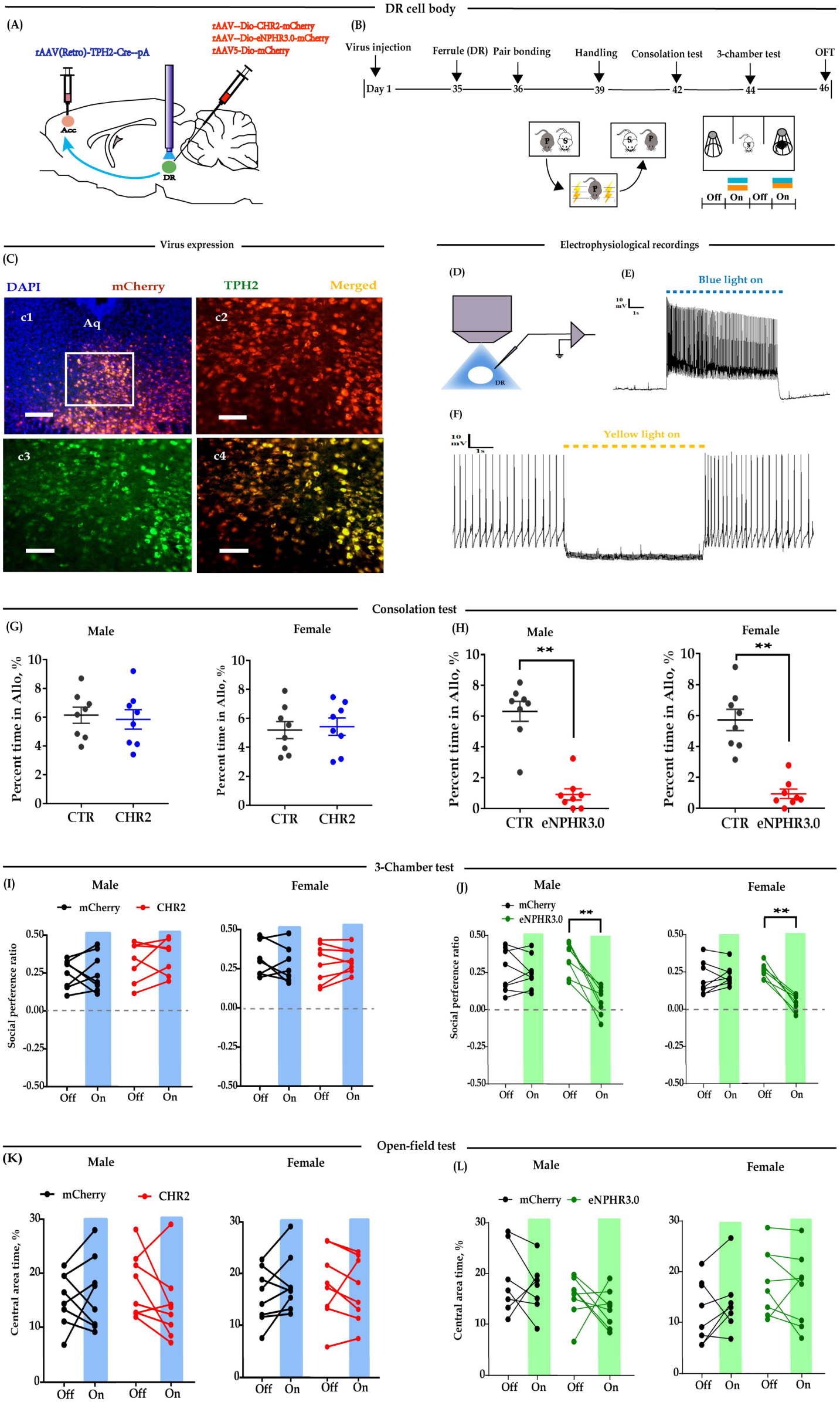
Optogenetic bidirectional modulation of 5-HT neuron in the DR in the DR-ACC neural circuit. (A): Schematic of optogenetic manipulation; (B): timeline of experiments; (C): immunohistological image showing virus expression in the DR (c1, × 100) and amplified images in the left box showing the mCherry, TPH2 and the colocaliztion of the two (‘c2-c4’, × 200); (D): electrophysiological recordings model; (E & F): representative traces from electrophysiological recordings showing photostimulation (E) and photoinhibition of a 5-HT neuron (F); (G-L): quantification of allogrooming time in the consolation test (G & H); social preference ratio in the three-chamber test (I & J) and time spent in the central area in the open-field test (K & L) during optogenetic modulations. Data are presented as mean ± SE, *n* = 7–8 in each group, \*\**P* < 0.01. For ‘G-H’, independent samples *t* tests along with Bayesian independent samples *t*-test; for ‘I-L’, two-way repeated measures ANOVA along with two-way repeated Bayesian measures ANOVA (light as within subject factor). For raw data in this figure, please refer to Figure 1—source data 1. For detailed quantifications, please refer to Supplementary 2. ACC: anterior cingulate cortex; CTR: control; DR: dorsal raphe nucleus; TPH2: tryptophan hydroxylase 2.

To test whether modulation of DR 5-HT neuron activity alters consolation and sociability, five weeks after the virus injection, optic fibers were implanted above the DR (Figure 1A-B). The virus expression sites and the optic fiber placement schematics are shown in Figure 1—figure supplement 3A. According to Burkett’s and our previous results (Burkett et al., 2016; Li et al., 2019), the time spent grooming their stressed partners (allogrooming) is an important indicator of consolation-like behaviors. We then exposed the subjects to their electric-shocked partners and recorded the time spent allogrooming, chasing (closely following) and selfgrooming (consolation test; for details, please refer to the Methods section). The three-chamber test was used to assay the sociability along with open-field anxiety and locomotion (Figure 1B). In the CHR2-expressing animals, there was no conclusive evidence that optogenetic activation of 5-HTergic neurons affected their time spent allogrooming, chasing and self-grooming in the 10-min consolation test (Figure 1G, Figure 1—figure supplement 4A-D), the social interaction ratio in the 3-chamber test (Figure 1I), and behavioral performance in the open-field test (Figure 1K, Figure 1—figure supplement 4E).

However, in the eNPHR3.0-expressing animals, there was extremely strong evidence that optogenetic inhibition of 5-HT neurons in the DR reduced the time spent allogrooming (Figure 1H) and chasing (Figure 1—figure supplement 4B), but had little effect on the control behavior self-grooming (Figure 1—figure supplement 4D). Following experiments indicated that the light-inhibition effects significantly reduced within 24 h (Figure 1—figure supplement 5). In the 3-chamber test, light inhibition also significantly reduced the social preference ratio (Figure 1J). In the open-field test, the results were inconclusive, which suggests that the data for the time spent in the central area and the total distance travelled are equally likely under H0 and H1 (Figure 1L, Figure 1—figure supplement 4F).

The above results indicate that although we have no conclusive results for optogenetic activation, optogenetic inhibition of 5-HT neurons within the DR→ACC 5-HTergic circuit impaired consolation-like behavior and sociability in mandarin voles.

### Optogenetic inhibition of ACC 5-HT terminals in the DR→ACC neural circuit similarly impaired consolation and reduced sociability

Modulation of 5-HT neurons may affect other neurons in the DR, and thus confound behavioral performance. In subsequent experiments, we placed optic fibers in the ACC and investigated whether the direct modulation of 5-HT terminals in this region would exert the same effects (Figure 2A). Similarly, we found that optogenetic activation of DR-ACC 5-HT terminals did not significantly alter behavioral performance in the consolation test (Figure 2C, Figure 2—figure supplement 1A, 1C) and open-field test (Figure 2G, Figure 2—figure supplement 1E). In the 3-chamber test, two-way repeated measures ANOVA revealed a moderate ‘light × group’ interaction in females (F_(1,14) =_ 6.297, *P* = 0.025, BF_incl_ = 4.186). Subsequent *post hoc* comparison results showed that light stimulation slightly increased the social preference ratio in CHR2-expressing females but not in mCherry-expressing females (ChR2: off *vs*. on, *P* = 0.033; mCherry: off *vs*. on, *P* = 0.975; Figure 2E). The ‘light × group’ interaction was not evidently observed in males (F_(1,14) =_ 3.556, *P* = 0.08, BF_incl_ = 1.239).

**Figure 2.**
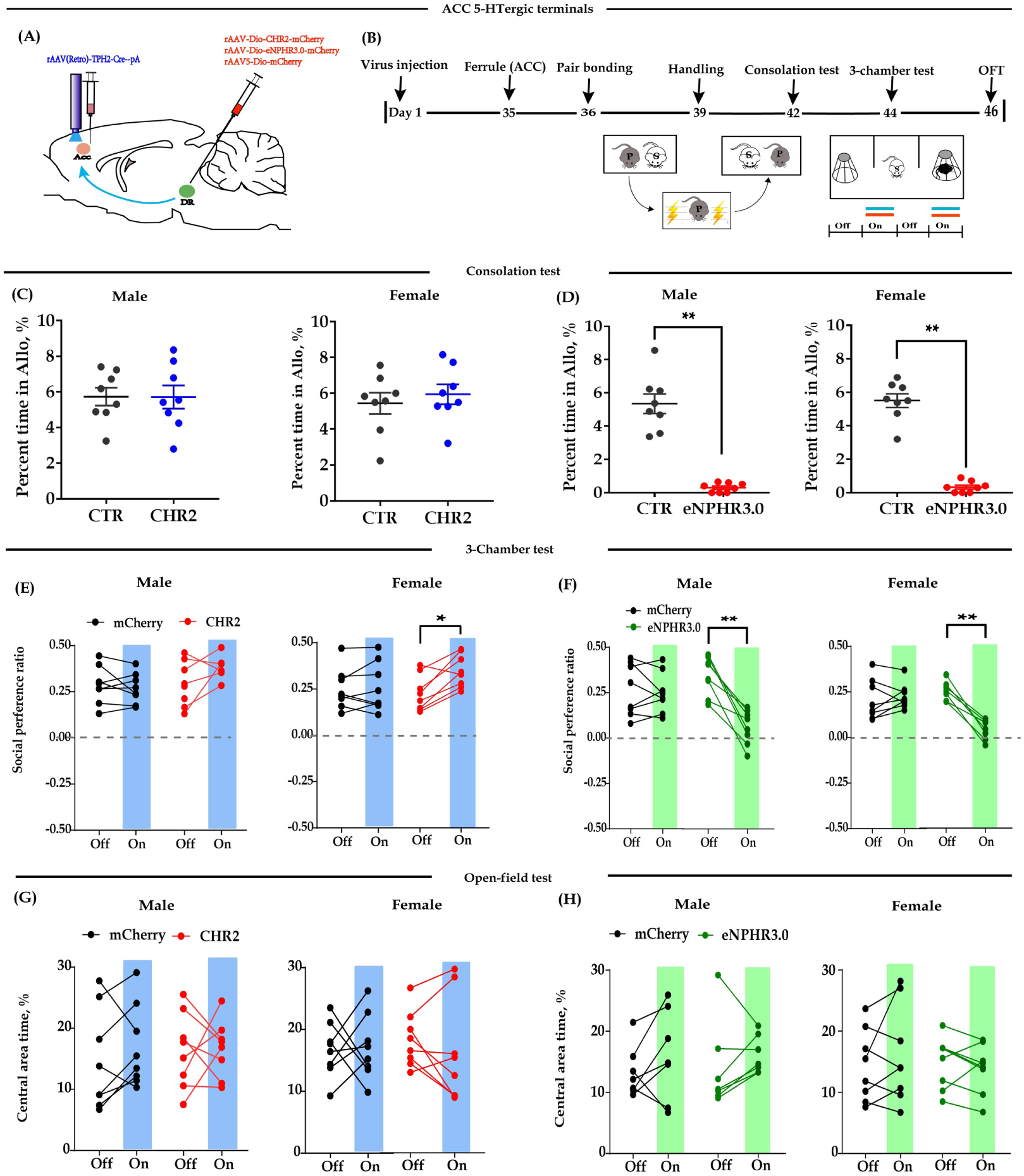
Optogenetic bidirectional modulation of 5-HT terminals within the ACC in the DR-ACC neural circuit. (A): Schematic of optogenetic manipulation; (B): timeline of experiments; (C-H): quantification of time spent in allogrooming in the consolation test (C & D), sociability in the three-chamber test (E & F) and time spent in the central area in the open-field test (G & H) during optogenetic modulations. Data are presented as mean ± SE, *n* = 7–8 in each group, \**P* < 0.05, \*\**P* < 0.01. For C-D, independent samples *t* tests along with Bayesian independent samples *t*-test; for E-H, two-way repeated measures ANOVA along with two-way repeated Bayesian measures ANOVA (light as within subject factor). For raw data in this figure, please refer to Figure 2—source data 1. For detailed quantifications, please refer to Supplementary 2. ACC: anterior cingulate cortex; DR: dorsal raphe nucleus; CTR: control.

Optogenetic inhibition of DR-ACC 5-HT terminals significantly reduced the time spent allogrooming and chasing in the consolation test (Figure 2D, Figure 2—figure supplement 1B) and the social preference ratio in the 3-chamber test (Figure 2F). Similarly, the inhibition had little effect on the control behavior of selfgrooming (Figure 2—figure supplement 1D) and behavioral performance in open-field test (Figure 2H, Figure 2—figure supplement 1F) and the inhibitory effect lasted no more than 24 h (Figure 2—figure supplement 2). The above results indicated that optogenetic modulation of ACC 5-HT terminals have nearly identical effects as modulation of DR 5-HT neurons.

### Chemogenetic inhibition of DR 5-HT neurons in the DR→ACC neural circuit impaired consolation and reduced sociability

To confirm the above optogenetic results over a longer time frame, we used a chemogenetic approach to selectively express ‘Gq-DREADD’ or ‘Gi-DREADD’ in DR 5-HT neurons by injecting AAV-DIO-hM4Dq-mCherry (Gq-DREADD) or AAV-DIO-hM4Di-mCherry (Gi-DREADD) into the DR and rAAV(Retro)-TPH2-Cre into the ACC (Figure 3A). Immunohistochemical staining revealed that more than 65% of TPH2 labeled neurons were infected by mCherry virus and more than 60% of mCherry cells coexpressed TPH2 (Figure 3—figure supplement 1). The virus expression sites schematics were shown in Figure 1—figure supplement 3B. To determine whether the ligand CNO can activate or inhibit DR 5-HT neurons, whole-cell current-clamp recordings were performed. The results showed that the addition of 10 μM CNO remarkably increased the number of action potentials in the Gq-DREADD-transfected neurons (Figure 3D). In contrast, CNO caused a significantly decrease in the number of spikes (Figure 3E) and increased the spike rheobase during current step injections in Gi-DREADD-transfected neurons (Figure 3F). These results indicate the specificity and viability of this virus strategy.

**Figure 3.**
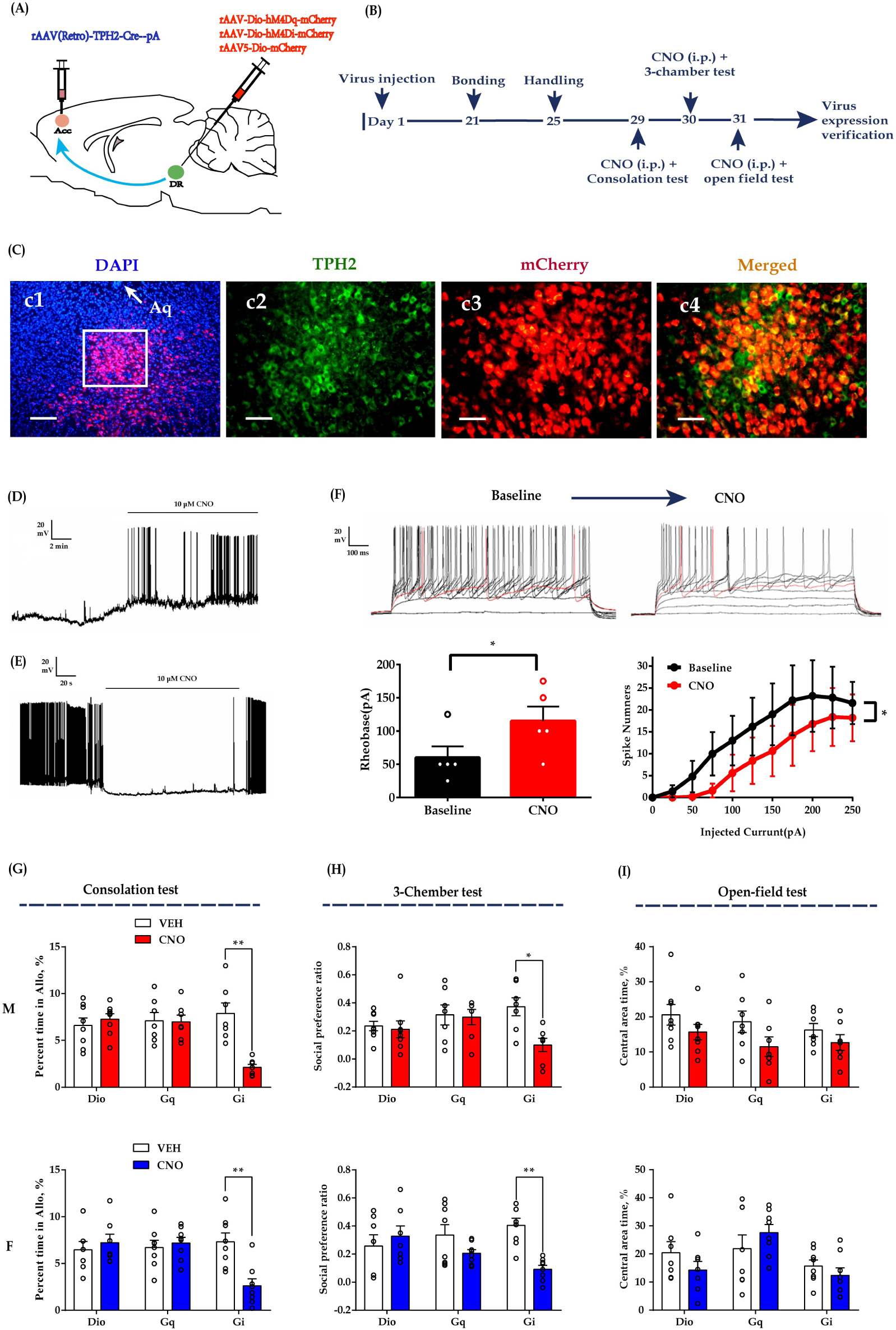
Chemogenetic modulation of DR 5-HT neuron activities in the DR-ACC neural circuit. (A): Schematic of chemogenetical manipulations; (B): timeline of experiments; (C): immunohistological image showing virus expression in the DR (c1, × 100) and amplified images in the left white box showing the mCherry, TPH2 and the colocaliztion of the two (c2-c4, × 200); (D): representative trace from a Gq-DREADD neuron; (E): representative trace from a Gi-DREADD-transfected neuron; (F): quantification of spike rheobase and spike numbers under current step injections in Gi-DREADD-transfected neurons (*n* = 5 neurons, \**P* < 0.05); (G-I): quantification of allogrooming time in the consolation test (G); social preference ratio in the three-chamber test (H) and time spent in the central area in the open-field test (I) in male (upper panels) and female (down panels) voles. Data are presented as mean ± SE, *n* = 7–8 in each group, \**P* < 0.05, \*\**P* < 0.01 compared with vehicle control. For ‘F’, one tailed paired *t* test along with one tailed Bayesian paired samples *t*-test; for ‘G-I’, two-way repeated measures ANOVA along with two-way repeated Bayesian measures ANOVA. For raw data in this figure, please refer to Figure 3—source data 1. For detailed quantifications, please refer to Supplementary 2. ACC: anterior cingulate cortex; DR: dorsal raphe nucleus; TPH2: tryptophan hydroxylase 2; M: male; F: female.

In subsequent behavioral studies, we found CNO (1 mg/kg, i.p.) treated Gi-DREADD-expressing voles showed reduced grooming toward their shocked partners in the consolation test (Figure 3G) and decreased sociability in the 3-chamber test (Figure 3H). The CNO treated males also showed a trend of spending less time in the open-field test, but the statistics were inconclusive (Figure 3I; treatment: F_(1,19)_ = 4.305, *P* = 0.052, BF(incl) = 4.438; Group: F_(2,19)_ = 2.064, *P* = 0.155, BF(incl) = 0.384; treatment × Group: F_(1,19)_ =0.164, *P* = 0.850, BF(incl) = 0.300). The treatment had no detectable effects on the total distance travelled in the open-field test (Figure 3—figure supplement 2C) and some control behaviors in the consolation test (Figure 3—figure supplement 2A&B). In Gq-DREADD and virus control subjects, CNO treatment had no detectable effects on the behavioral performance in all the test (Figure 3).

Based on the results of the chemogenetic and optogenetic experiments described above, we concluded that inhibition of DR-ACC 5-HTergic circuit activity was sufficient to impair consolation and sociability in mandarin voles.

### DR 5-HT neuron activity increased during allogrooming and social approaching

If the DR-ACC 5-HTergic circuit is involved in consolation and sociability, ACC projecting DR 5-HT neurons may change their activity during the corresponding behaviors. To verify this idea, we performed photometric recording in freely moving animals by injecting Cre-inducible AAV-DIO-GCaMp6, a genetically encoded fluorescent Ca^2+^ sensor, into the DR and rAAV(Retro)-TPH2-Cre into the ACC (Figure 4A). Ten days later, fibers were implanted above the DR injection sites. Post hoc histological analysis revealed that more than 80% GCaMp6^+^ cells overlapped with TPH2^+^ expressing cells (Figure 4—figure supplement 1). The virus expression sites and the optic fiber placements schematics were shown in Figure 1—figure supplement 3C.

During the consolation test, we found that the activity of DR 5-HT neurons was reliably increased during allogrooming, social approaching and sniffing in both sexes (Figure 4E1-E2, F1-F2 and G1-G2). DR 5-HT neurons showed the highest response during allogrooming which may indicate the specific role of 5-HT in consolation-like behaviors (Figure 4—figure supplement 2A-2B). Interesting, the increased Ca^2+^ in DR 5-HT neurons often preceded these behaviors which may indicate the motivational function of 5HT (Arakawa, 2020; Yagishita, 2020). When aligning the fluorescence changes to the end of these social bouts, we found that the activity of 5-HT neurons decreased accordingly before withdrawal from allogrooming and approaching (Figure 4—figure supplement 3A-3B). Withdrawing from sniffing also showed a similar trend but was not statistically significant (Figure 4—figure supplement 3C; pre *vs*. post: male: t_(5)_ = 1.611, *P* = 0.084, BF+0 = 1.589 with median posterior δ = 0.528, 95%CI = [0.041, 1.363]; female: t_(5)_ = 1.724, *P* = 0.073, BF+0 = 1.778 with median posterior δ = 0.557, 95%CI = [0.046, 1.415]). The increased activity of DR 5-HT neurons did not simply reflect movement initiation as no fluorescence changes were observed when aligning to running and selfgrooming (Figure 4H1-H2, I1-I2). Furthermore, no significant changes in fluorescence signals were detected in DR 5-HT neurons of GFP-expressing control voles during all the behaviors, confirming that the GCaMp6 signals genuinely indicated neuronal activity and did not simply reflecting motion artifacts (Figure 4E3-E4, F3-F4, G3-G4, H3-H4 and I3-I4).

**Figure 4.**
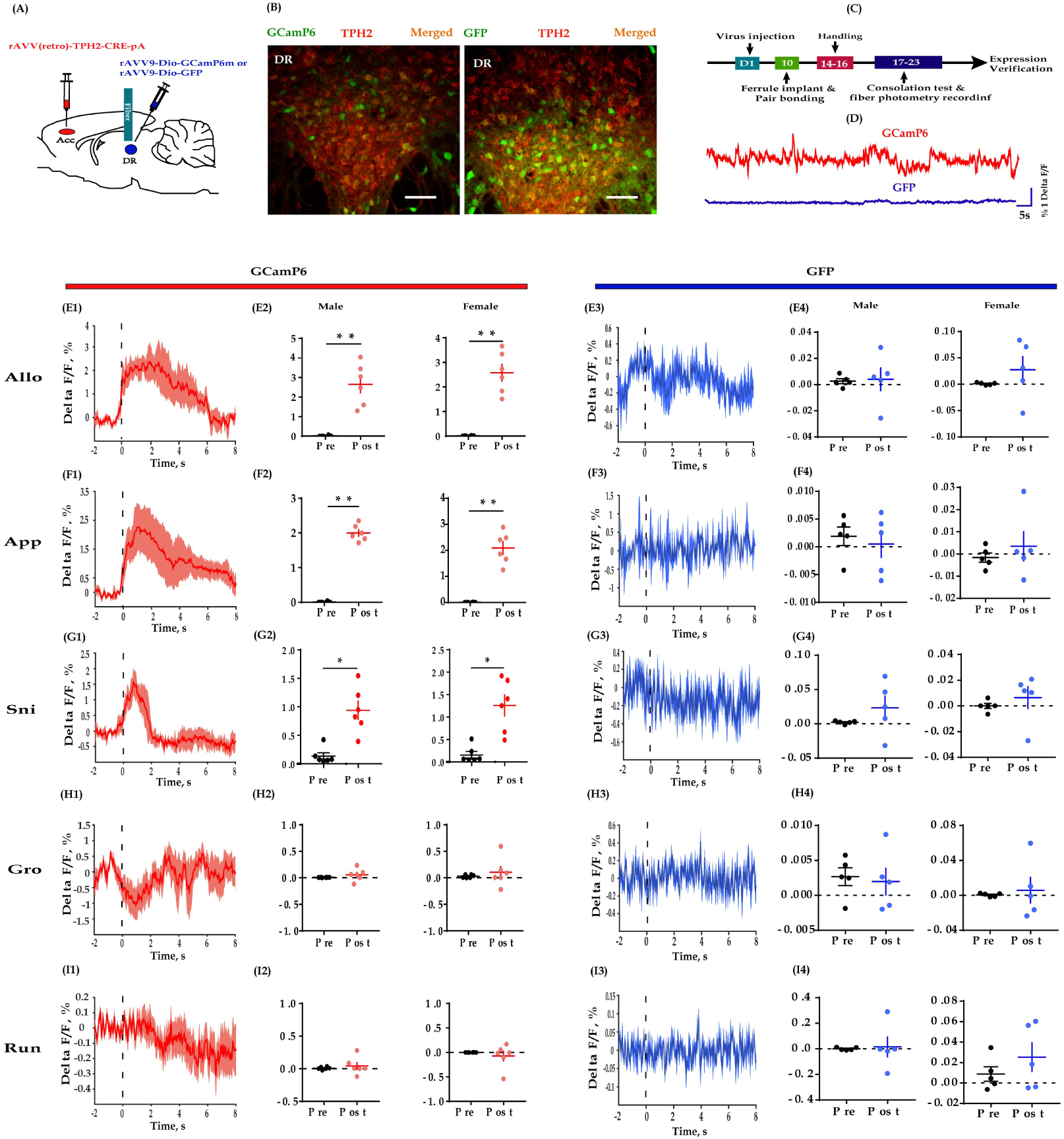
Fiber photometry recording DR 5-HT neural dynamics during the consolation test. (A): Schematic diagrams depicting the virus injection and recording sites; (B): histology showing the expression of GCaMP6 (left) and GFP control (right) in the DR (× 200); (C): experimental timeline for photometry experiments; (D): representative fluorescence changes of GCaMP6 (red line) and GFP (blue line) during photometry recordings; (E1-I1): representative peri-event plot of GCaMP6 fluorescence signals aligned to onsets of various behavior (for all peri-event plots, the red line denotes the mean signals of 4-6 bouts of behaviors, whereas the red shaded region denotes the SE); (E2-I2): quantification of change in GCaMP6 fluorescence signals before and after the events (*n* =6 in each group); (E3-I3): representative peri-event plot of GFP signals aligned to onsets of various behavioral events (for all peri-event plots, the blue line denotes the mean signals of 4-6 bouts of behaviors, whereas the blue shaded region denotes the SE); (E4-I4): quantification of change in GFP fluorescence signals before and after the events (*n* = 5 in each group). Data are presented as mean ± SE, \**P* < 0.05, \*\**P* < 0.01, Paired samples *t*-test along with Bayesian paired samples *t*-test. For raw data in this figure, please refer to Figure 4—source data 1. For detailed quantifications, please refer to Supplementary 2. ACC: anterior cingulate cortex; TPH2: tryptophan hydroxylase 2; Allo: allogrooming; Sni: sniffing; App: approaching; Gro: selfgrooming; Run: running.

The above results provide direct evidence that DR 5-HT neurons within the DR-ACC 5-HTergic circuit are recruited in consolation-like behavior and sociability in mandarin voles.

### ACC 5-HT release increased during allogrooming, social approach and sniffing

To further establish the potential relevance of 5-HT in consolation-like behaviors, we directly detected the dynamics of 5-HT within the ACC using a newly developed GPCR-activation-based 5-HT sensor along with photometric recording (Wan, *et al* 2020). To do this, we injected AAV encoding the fluorescent 5-HT sensor (AAV-5HTsensor 2.1) unilaterally into the ACC and implanted optic fibers above the injection site ten days later (Figure 5A). The virus expression sites and the optic fiber placements schematics were shown in Figure 1—figure supplement 3D. Control animals were injected with AAVs encoding eGFP. Post hoc histological verification indicated expression of both the 5-HT sensor and eGFP within the ACC (Figure 5B).

**Figure 5.**
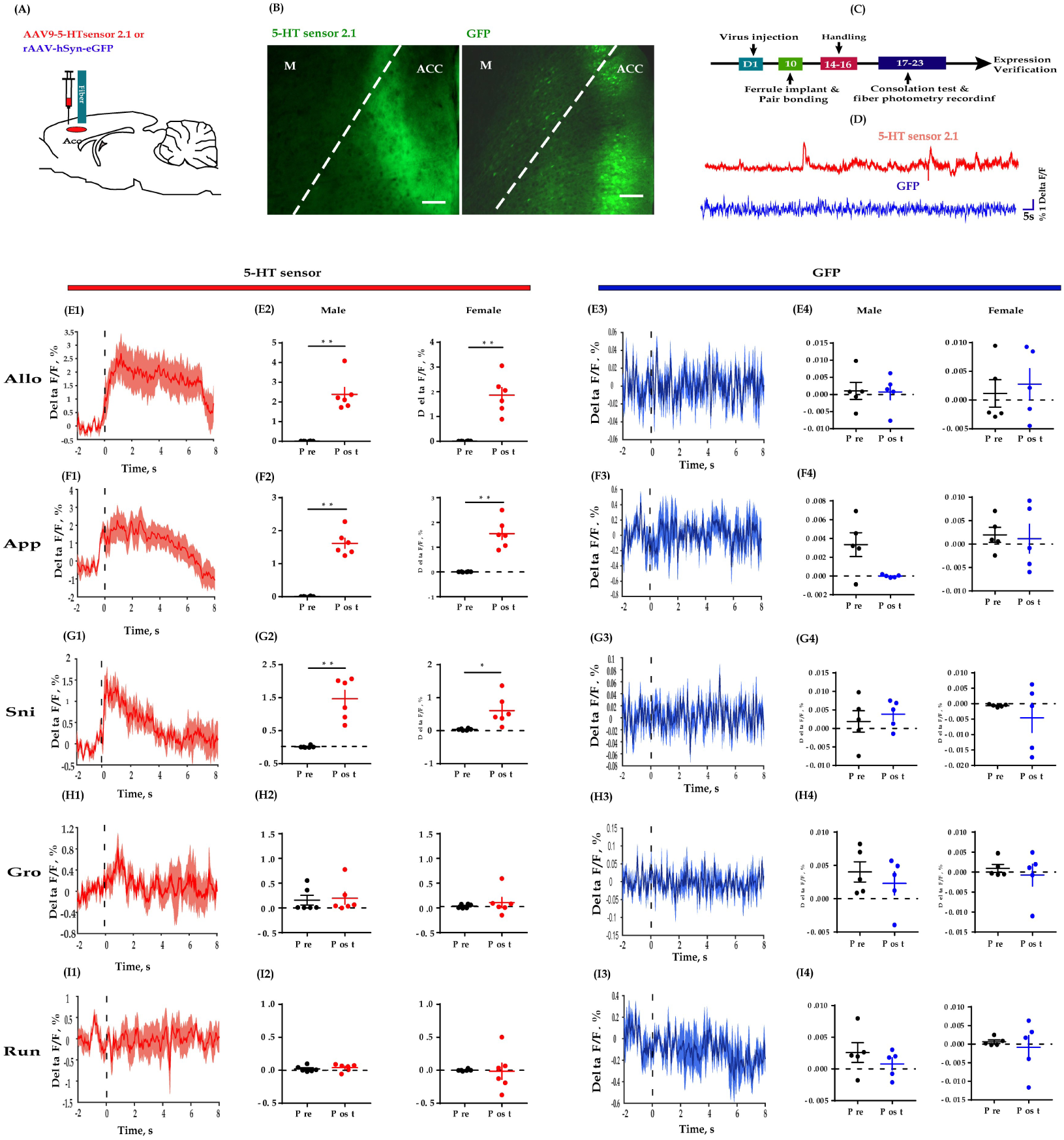
Fiber photometry recording dynamics of 5-HT within the ACC during the consolation test. (A): Schematic diagrams depicting the virus injection and recording sites; (B): histology showing the expression of 5-HT sensor (left) and GFP control (right) within the ACC (× 200); (C): experimental timeline for photometry experiments; (D): representative fluorescence changes of 5-HT sensor (red line) and GFP (blue line) during photometry recordings; (E1-I1): representative peri-event plot of 5-HT fluorescence signals aligned to onsets of various behavior (for all peri-event plots, the red line denotes the mean signals of 4-6 bouts of behaviors, whereas the red shaded region denotes the SE); (E2-I2): quantification of change in 5-HT fluorescence signals before and after the events (*n* = 6 in each group); (E3-I3): representative peri-event plot of GFP signals aligned to onsets of various behavioral events (for all peri-event plots, the blue line denotes the mean signals of 4-6 bouts of behaviors, whereas the blue shaded region denotes the SE); (E4-I4): quantification of change in GFP fluorescence signals before and after the events (*n* = 5 in each group). Data are presented as mean ± SE, \**P* < 0.05, \*\**P* < 0.01; paired samples *t*-test along with Bayesian paired samples *t*-test. For raw data in this figure, please refer to Figure 5—source data 1. For detailed quantifications, please refer to Supplementary 2. ACC: anterior cingulate cortex; M: motor cortex; TPH2: tryptophan hydroxylase 2; Allo: allogrooming; Sni: sniffing; App: approaching; Gro: selfgrooming; Run: running.

Next, we measured the dynamics of endogenous 5-HT during the consolation test. Consistent with the above fiber photometry of Ca^2+^ signals in DR 5-HT neurons, we found reliably increased fluctuations in fluorescence within the ACC in conjunction with allogrooming, social approaching and sniffing, but not observed with self-grooming and running (Figure 5, left columns). Similarly, the peek 5-HT signals during allogrooming were larger than during the other behaviors (Figure 4—figure supplement 2C-2D), and the release of 5-HT significantly dropped before the end of allogrooming, approaching and sniffing (Figure 5—figure supplement 1). In control animals that expressed GFP in the ACC, we observed little fluorescence change during any of these behaviors (Figure 5E3-E4, F3-F4, G3-G4, H3-H4 and I3-I4).

To validate the selectivity of this 5-HT sensor in mandarin voles, in another set of animals expressing the 5-HT sensor, a 5-HT receptor antagonist metergoline (Met) was injected (i.p., 4 mg/kg) before conducting the fiber photometry experiments. We found that treating voles with Met completely blocked the fluorescence changes of the 5-HT sensor during allogrooming (Figure 5—figure supplement 2), validating the viability and specificity of this 5-HT sensors.

The above results provide further evidence that 5-HT released within the ACC may play an important role in consolation-like behavior and sociability in mandarin voles.

### Serotonin in the ACC mediated consolation-like behaviors through 5-HT1A receptors

The most abundant 5-HT receptors expressed in the mPFC are 5-HT1AR and 5-HT2AR (Carhart-Harris and Nutt, 2017). Increasing evidence indicates that these two types of receptors exert opposite effects within the mPFC (Celada et al., 2013; Puig and Gulledge, 2011). In our previous study, we indicated that ACC administration of a 5HT1AR agonist (8-OH-DPAT) rescued the impaired consolation and sociability induced by social defeat (Li et al., 2020). To further examine which receptor mediates consolation-like behaviors within the ACC, we directly infused 8-OH-DPAT (200 nL/side; 1.5 mg/mL) or a 5-HT2AR antagonist (MDL 100907, 200 nL/side; 1 mg/mL) into the ACC along with chemogenetic inhibition of DR 5-HT neurons before conducting behavioral assays (Figure 6A & B). Not surprisingly, CNO elicited deficits in allogrooming and sociability (Figure 6C, D, F, G: left two bars). Pretreatment with 8-OH-DPAT significantly reduced the impact of CNO (Figure 6C, D: right two bars; see also the comparison results of ‘Vehicle+Saline - Vehicle+CNO’ *vs*. ‘8-OH-DPAT+Saline - 8-OH-DPAT+CNO’; Figure 6—figure supplement 1A, B), whereas MDL 100907 had no such effect (Figure 5F, G: right two bars; Figure 6—figure supplement 1C, D). Neither the drugs nor viruses had any detectable effect on the control behavior of self-grooming or behavioral performance in the open-field test (Figure 6E-H, Figure 6—figure supplement 2).

**Figure 6.**
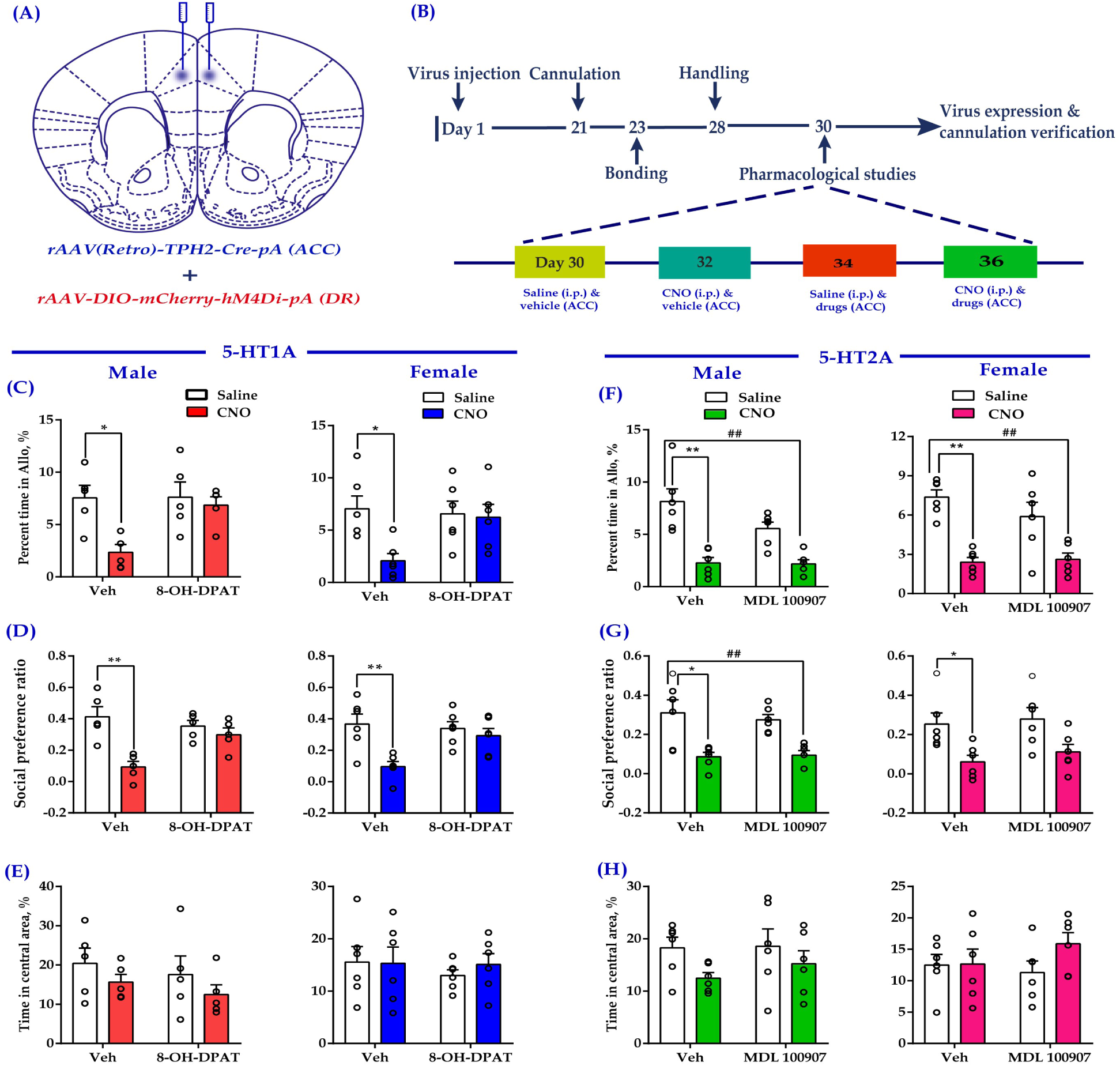
Intra-ACC injection of 8-OH-DPAT reduced sociability deficits induced by chemogenetic inhibition of DR 5-HT neurons in the DR→ACC neural circuit. (A): Schematic representation of ACC infusion sites and virus strategy; (B): timeline of experimental design; (C-E): effect of a 5-HT1AR agonist 8-OH-DPAT on allogrooming time in the consolation test (C), social preference ratio in the three-chamber test (D), and time spent in the central area in the open-field test (E) in male (left panels) and female (right panels) voles; (F-H): effect of a 5-HT2AR antagonist (MDL 100907) on allogrooming time in the consolation test (F), social preference ratio in the three-chamber test (G) and time spent in the central area in the open-field test (H) in male (left panels) and female (right panels) voles. Data are presented as mean ± SE, *n* = 5–6 in each group; two-way ANOVA along with two-way Bayesian ANOVA; \**P* < 0.05, \*\**P* < 0.01 for ‘vehicle + saline’ *vs.* ‘vehicle + CNO’; ^##^*P* < 0.01 for ‘MDL 100907 + CNO’ *vs.* ‘vehicle + saline’. For raw data in this figure, please refer to Figure 6—source data 1. For detailed quantifications, please refer to Supplementary 2. Anta: antangonist; ACC: anterior cingulate cortex; DR: dorsal raphe nucleus.

## DISCUSSION

In the present study, we demonstrated a crucial role for the DR→ACC 5-HTergic neural circuit in the regulation of consolation-like behaviors and sociability in mandarin voles for the first time. Our major findings are listed below. First, inhibition of DR 5-HT neurons or their terminals in the ACC decreased allogrooming behavior and reduced sociability. Second, DR 5-HT neuron activity and ACC 5-HT release increased during allogrooming, social approaching and sniffing. Third, direct activation of ACC 5-HT1A receptors was sufficient to ameliorate deficits in consolation and sociability induced by chemogenetic inhibition of DR 5-HT neurons.

We found optogenetic inhibition of DR 5-HT neurons (Figure 1) or ACC 5-HT terminals (Figure 2) significantly decreased intimate behaviors (allogrooming and chasing) toward their distressed partners and reduced sociability. Chemogenetic inhibition of DR 5-HT neurons produced similar results (Figure 3). Consistent with our results, Walsh et al found that optogenetic inhibition of DR 5-HT neurons reduced social interactions in the three-chamber test (Walsh et al., 2018). As consolation is in general a pro-social behavior, it is difficult to determine whether the reduced allogrooming is due to an overall decrease in sociability. However, in the fiber photometry studies, we found both DR 5-HT neurons and ACC 5-HT release showed the highest responses during allogrooming, which may indicate the specific role of 5-HT in consolation (Figure 4—figure supplement 2).

Another open question is how this process occurs, namely, what are the neural mechanisms underlying this process? The neocortical excitatory/inhibitory (E/I) balance hypothesis may help address this question. The hypothesis indicates that an increase in the cortical cellular E/I balance, for example through increased activity in excitatory neurons or a reduction in inhibitory neuron function, is the common etiology and final pathway for some psychiatric diseases, such as autism and schizophrenia (Bozzi et al., 2018; Vattikuti and Chow, 2010). This hypothesis has recently been verified in mice, as optogenetic excitation of glutamatergic neurons in the mPFC elicits a profound impairment in sociability, while compensatory excitation of inhibitory neurons in this region partially rescues the social deficits caused by an increase in the E/I balance (Yizhar et al., 2011). 5-HT tends to inhibit prefrontal pyramidal activity (Puig and Gulledge, 2011; Tian et al., 2017). For example, according to Puig et al., electrical stimulation of the DR inhibits approximately two-thirds of pyramidal neurons in the mPFC (Puig et al., 2005). Therefore, a reasonable hypothesis is that inhibition of the DR-ACC 5-HTergic neural circuit may increase the E/I balance, which ultimately leads to abnormal social behaviors in mandarin voles. Clearly, this hypothesis should be verified in electrophysiological studies in the future.

In contrast to inhibition manipulations, we found activation of the DR-ACC 5-HTergic neural circuit did not elicit corresponding increases in allogrooming and sociability (Figures 1-3). One possible explanation is that there are various 5-HT receptors expressed in the ACC and some have opposite functional effects (Puig and Gulledge, 2011; Tian et al., 2017). For example, 5-HT1AR coupled to the Gi family of G proteins induces hyperpolarization of pyramidal neurons, whereas 5-HT2AR coupled to Gq proteins induces depolarization in the same neurons (Carhart-Harris and Nutt, 2017). In addition, these two types of receptors are expressed in both pyramidal and GABAergic neurons of the mPFC (Santana et al., 2004). Therefore, the net behavioral effects of 5-HT release must result from the combined effects of all these receptors. The other possible explanation is a ceiling effect, which means that the enhancement of consolation and sociability are difficult to accomplish in these normal animals. This may occur in situations where levels of consolation and sociability are low such as in the stressed subjects in our previous study (Li et al., 2020) or genetically less sociable subjects as in Walsh’s study (Walsh et al., 2018). However, in a previous study by Kim et al., the direct administration of 5-HT into the ACC impaired observational fear learning, which is an indicator of emotional contagion (Kim et al., 2014). Species differences (mice *vs.* voles), methodological differences (pharmacology *vs*. optogenetics and chemogenetics) or different behavioral indicators (consolation *vs*. emotional contagion) may account for the discrepancies. Furthermore, optogenetic or chemogenetic activation of DR 5-HT neurons is also distinct from pharmacological intervention with MDMA, which robustly induces 5-HT releases in the whole brain and enhances closeness and empathy in its users in human studies (Carlyle et al., 2019; Heifets and Malenka, 2016). Nevertheless, at least some effects of activation were visible. For example, optogenetic activation of ACC 5-HT terminals increased sociability in CHR2-expressing females (Figure 2E). Clearly, the effects of activation of the DR-ACC 5-HTergic neural circuit on sociability and empathy still require further in-depth study.

Our results show that neither activation nor inhibition the DR-ACC 5-HTergic neural circuit exerts any significant effects on some control behaviors (Figure 1—figure supplement 4; Figure 2—figure supplement 1; Figure 3—figure supplement 2), i.e., self-grooming in the consolation test or behavioral performance in the open-field test, which is consistent with pervious results (for self-grooming, please refer to (Correia et al., 2017); for the open-field test, please refer to (Walsh et al., 2018)). However, inconsistent results have also been reported. For example, Ohmura et al. found that optogenetic activation of 5-HT neurons in the median raphe nucleus enhanced anxiety-like behaviors in mice (Ohmura et al., 2014). In the study by Correia et al., activation of 5-HTergic neurons in the DR affected behavior in the open-field test, but not anxiety (Correia et al., 2017). Different stimulation protocols or manipulations of different groups of 5-HT neurons may account for these discrepancies.

Our results obtained using *in vivo* fiber photometry indicated an increase in DR 5-HT neuron activity during allogrooming, sniffing and social approaching (Figure 4). This result is consistent with previous studies showing increases in the activity of 5-HTergic neurons in the DR during nonaggressive social interactions (Li et al., 2016; Walsh et al., 2018). Furthermore, using the highly sensitive 5-HT fluorescent sensors developed by Wan et al., (Wan et al, 2020), we provided the first evidence that allogrooming, sniffing and social approaching elicit robust 5-HT release in the ACC (Figure 5). Self-grooming and running did not induce robust 5-HT release within the ACC, which may indicate the behavioral relevance of the 5-HT sensors. Interestingly, we found that the fluorescence changes in GCamP6 and 5-HT sensors usually precede allogrooming and social approaching and decreased accordingly before withdrawing from them (Figure 4—figure supplement 3, Figure 5—figure supplement 1). These results indicate that 5-HT may be involved in some motivational aspects of these social behaviors, which is consistent with previous findings (Arakawa, 2020; Yagishita, 2020).

Although both 5-HT1AR and 5-HT2AR are expressed at high levels in the mPFC and work together to modulate cortical network activity (Celada et al., 2013), we found only the 5-HT1AR receptor agonist significantly reduced the consolation and sociability deficits induced by chemogenetic inhibition of 5-HTergic neurons in the DR (Figure 6). This result is consistent with our previous finding that the administration of WAY-100635 (a 5-HT1AR antagonist) into the ACC attenuates consolation and sociability in mandarin voles (Li et al., 2020). 5-HT1AR within the mPFC has frequently been reported to exert antidepressant and anxiolytic effects (Artigas, 2013; Fukumoto et al., 2020; Fukumoto et al., 2014; Wang et al., 2019), but our findings clearly indicate that this type of 5-HT receptor within the ACC is also involved in regulating some social behaviors. Clearly, this effect should be verified in many other species in future studies. Administration of the 5-HT2AR antagonist into the ACC did not exert an obvious effect on the social deficits induced by the chemogenetic inhibition of DR 5-HT neurons (Figure 5F-H). Thus, 5-HT2AR within the ACC may not play a major role in consolation and sociability. However, this conclusion should be interpreted very cautiously because 5-HT2AR expression within the mPFC shows clear rostral-to-caudal gradients in mice (Weber and Andrade, 2010) and we were unable to determine its functional role in this dimension in the present study. Furthermore, the effects of 5-HT in the ACC on consolation and sociability probably involve other subtypes of 5-HT receptors, such as 5-HT1BR and 5-HT3R, which require further examination.

In conclusion, our findings establish the importance of the DR-ACC 5-HTergic neural circuit in consolation-like behaviors and sociability in mandarin voles. One major limitation of our study is that although our virus strategies were generally successful, a few of the infected neurons were indeed TPH2-negative, which may challenge the specificity of our manipulations and confound our results. However, the fiber photometry and pharmacological studies mitigate these concerns. Considering the widespread innervation of 5-HT terminals and abundant distribution of 5-HT receptors in the whole brain, detailed knowledge of cellular and circuit mechanisms of the 5-HTergic system, will not only improve our understanding of its complicated features and functions but also have implications for the development of novel therapies for the treatment of prevalent neuropsychiatric disorders, such as depression, autism, and schizophrenia. Additionally, although both male and female subjects were included in this study, sexually dimorphic effects were rarely observed. This finding may provide additional evidence of cooperative evolution to adapt to environmental challenges, particularly in species that adapt monogamous relationships and require cooperative breeding.

## METHODS AND MATERIALS

### Key Resources Table

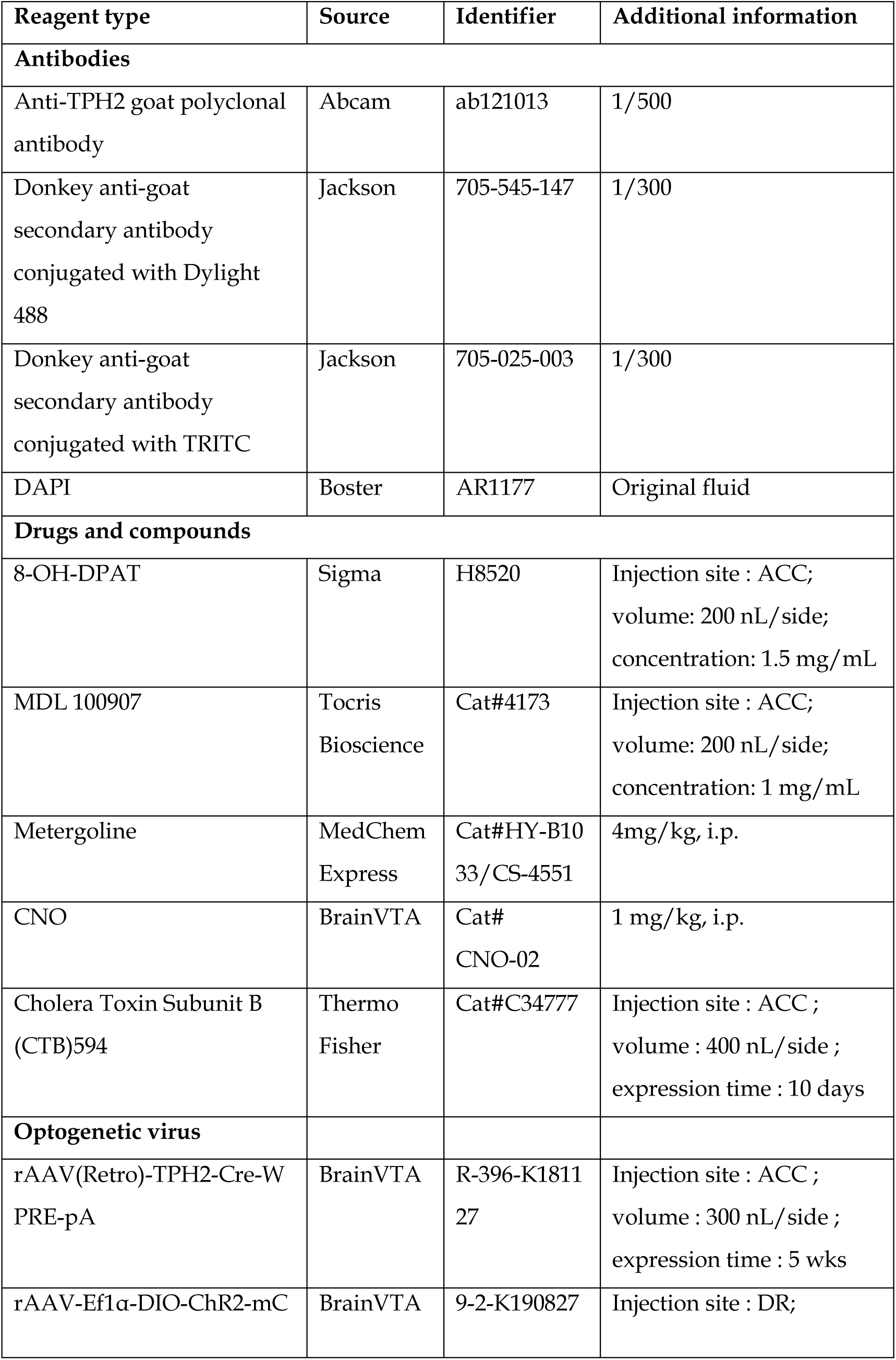
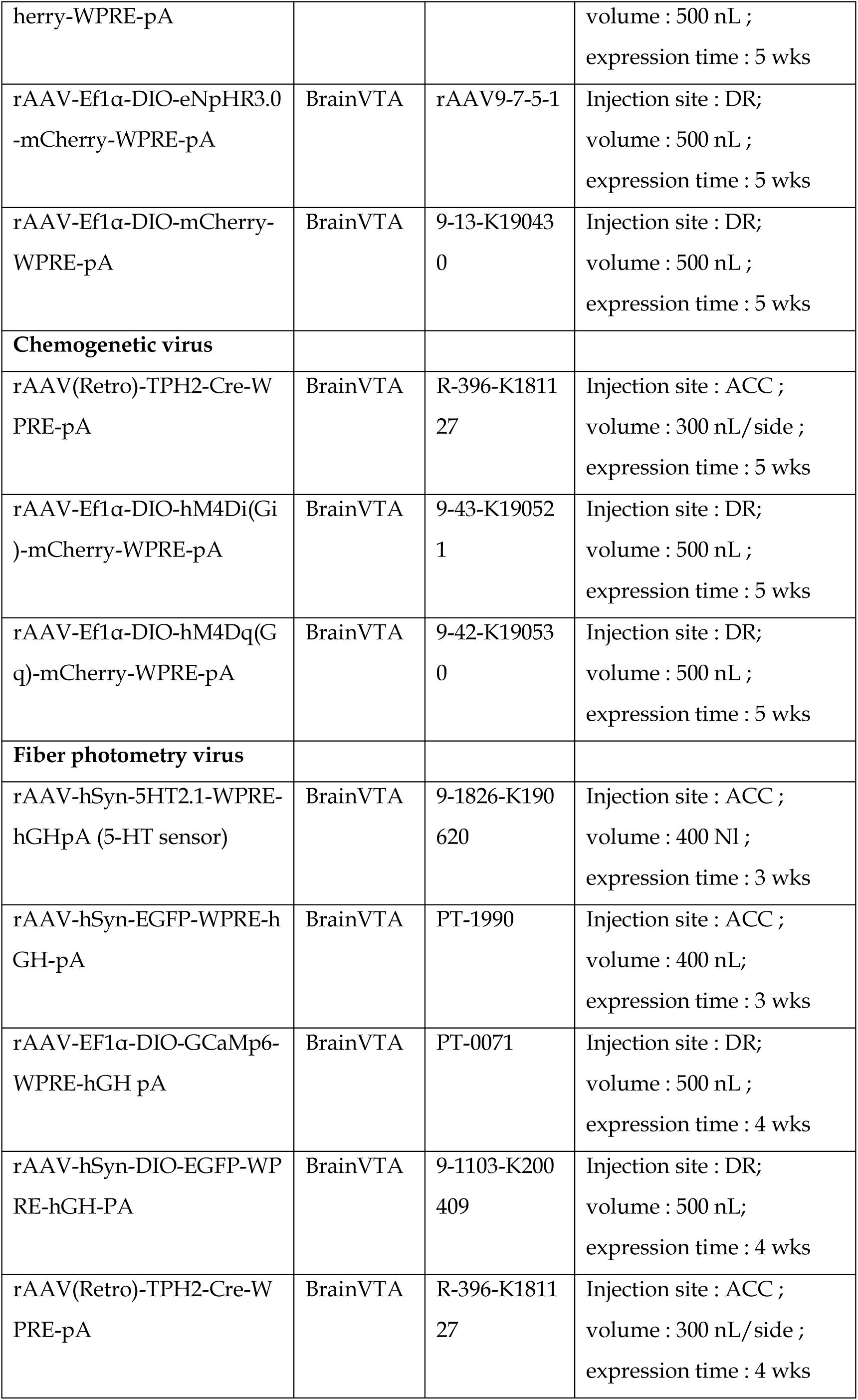

### Animals

The mandarin voles used in this study were laboratory bred strains (F2-F3) whose ancestors derived from a wild population from Lingbao city (Henan, China). The voles were weaned on postnatal day 21, socially housed in same-sex in each polycarbonate cage (44×22×16 cm) and housed on a 12-h light/dark cycle with food and water ad libitum. Voles used for experiments were about 70-90 days old at the time. All breeding, housing, and experimental procedures were in accordance with Chinese guidelines for the care and use of laboratory animals and were approved by the Animal Care and Use Committee of Shaanxi Normal University.

### Stereotaxic surgery and virus infusions

The kind of virus, total injection volumes and the expression time were listed in Key Resources Table. For surgery, voles (about 50 days) were anesthetized with 1.5%-3.0% isoflurane and placed in a stereotaxic instrument. Next, thirty-three-gauge syringe needles (Hamilton) were used for virus delivery. The injection rates were set at 50 nL/min. After each injection, the needle was left in the brain for another 5 min before being slowly withdrawn in order to prevent the virus from leaking out. The bregma coordinates for the virus injection were as follows: ACC: A/P: 1.6, M/L: 0.5, D/V: 1.6; DR: AP: −4.3; DV: −3.3, ML: +1.2, with a 20 angle toward the midline in the coronal plane.

### Microinjection and drugs

The 5-HT1AR agonist 8-OH-DPAT was prepared in saline with a final concentration of 1.5 mg/mL. The 5-HT2AR antagonist MDL 100907 was prepared in 0.01 M PBS (adjust PH value with 0.1 M HCl to 6.4) with a final concentration of 1 mg/mL. All the microinjections were administered 30 min before the behavioral test. The speed of injection was 0.1 μL/min, and the total injection volume were 0.2 µL per side for all the drugs. The injector tips remained in situ for another 2 min for drug diffusions. The dose and timing of drug administration were chosen based on previous studies with 8-OH-DPAT (Cooper et al., 2008; Li et al., 2020) and MDL 100907 (Ishii et al., 2015; Pockros et al., 2011), which adjusted according to preliminary studies. In chemogenetic studies, CNO (1 mg/kg) was dissolved in saline and delivered intraperitoneally 30 min before the behavioral test.

### Optogenetic studies

For optogenetic activation, ferrules were connected (by patch cords) to a 473 nm laser diode through a FC/PC adaptor and a fiber optic rotary joint. The output parameters were: 10 ms, 20 Hz, 8 s on and 2 s off cycle, ~10 mW for terminal stimulation, ~5 mW for somatic stimulation. For optogenetic inhibition, ferrules were connected to a 593 nm laser diode. The output parameters were: 10 ms, 20 Hz, constant, ~10 mW for both terminal and somatic inhibition. The optogenetic stimulation parameters were chosen and slightly modified based on previous studies (Garcia-Garcia et al., 2018; Walsh et al., 2018; Zhao et al., 2011)

### Behavioral assays

Generally, all the behavioral experiments were performed under dim light during the dark phase of the light-dark cycle. In all tests, all groups of experimental voles were randomly selected and the observers were blinded to the treatments. During the test, if the subject (not the stimulus voles) showed very rare or no movements, we think these individuals did not show normal activity, and their data are not appropriate to include in the following analysis. For optogenetic and fiber photometry studies, all subjects were allowed to habituate to patch cords for at least three days and allowed 30 min acclimation to the connection before the experiment started.

#### Consolation test

The consolation test was performed as previously described (Burkett et al., 2016). Briefly, five days before the experiment, age-matched adult male and female voles were cohoused together to promote the formation of a pair bond (Yu et al., 2012). On the testing day, the subjects’ partners were gently transferred in a cup to a sound-proof electric shock chamber and then subjected to 10 rounds of moderately strong foot shocks (3 s, 0.8 mA, 2 min intertrial intervals). At the same time, the test voles were left undisturbed in their home cages. After the separation, the pairs were reunited, and the behavior of the subjects was recorded using a video camera for 10 min in the test room. The digital videos were viewed and quantified using J Watcher software (http://www.jwatcher.ucla.edu/). According to previous studies (Burkett et al., 2016; Li et al., 2019), allogrooming is designated as behavioral indicators of consolation, which defined as head contact with the body or head of their partner, accompanied by a rhythmic head movement. The duration of chasing (following the shocked partners) and selfgrooming were also collected and analyzed.

For optogenetic tests, it designed as a between-subjects test in which both groups (mCherry or opsin) received laser stimulation. The lasting effects of optogenetic inhibition were measured 24 h after the original test, i.e., consolation test was conducted once again without light stimulation in eNPHR 3.0 animals one day later.

#### Three-chamber test

The three-chamber test was used to assess the sociability of a subject (Horie et al., 2019; Walsh et al., 2018). The apparatus consisted of a rectangular box with three separate chambers (20 × 40 × 20 cm each). One side of each chamber contained a circular metal wire cage (stimulus animal cage, 11 cm high and 9 cm in diameter). One day before the test, all subjects were habituated to the arena for 5 min and an age- and sex-matched unfamiliar stimulus voles also habituated to the wire cages for 5 min. On the testing day, the test vole was placed in the center chamber and a ‘stimulus’ vole (housed in a same room with the test voles) was randomly placed into one of the wire cages. After 2 min, the partitioning walls between the chambers were removed and the test voles were allowed to explore freely for a 10 min session. The time spent exploring each chamber was automatically recorded using a video tracking system (Shanghai Xinruan, China). Sociability was calculated as follows: (time spent in the stimulus vole side − time spent in the empty side)/(time spent in the stimulus vole side + time spent in the empty side).

For optogenetic tests, voles received four epochs of light beginning with a light OFF baseline epoch (OFF-ON-OFF-ON). Each epoch lasted for 5 min and the total duration was 20 min.

#### Open-field test

The anxiety level and locomotor function of the subjects were assessed by using the open-field test (Flanigan et al., 2020; Walsh et al., 2018). Briefly, a square open field (50 cm × 50 cm × 25 cm) was virtually subdivided into 16 even square. The four central squares were designated as central area. At the beginning of the test, the test vole was placed into the center area facing away from the experimenter. Behavior was recorded for 5 min. Outcome measures were distance traveled, frequency entries and time spent in the central area.

For optogenetic tests, voles were tested twice in the same day in both light-off and light-on conditions with at least 2 h between sessions (within-subjects design).

### Fiber photometry

The fiber photometry recording set-up (ThinkerTech, Nanjing, China) was generated and used as previously described (Feng et al., 2019; Yuan et al., 2019). Briefly, the emission light was generated by a 480 LED, reflected with a dichroic mirror and delivered to the brain in order to excite the GCaMP6m or the 5-HT sensor. The emission light passed through another band pass filter, into a CMOS detector (Thorlabs, Inc. DCC3240M) and finally recorded by a Labview program (TDMSViewer, ThinkerTech, Nanjing, China).

On the test day, voles were mildly anesthetized with isoflurane and connected to a muti-mode optic fiber patch cord (ThinkerTech Nanjing Bioscience®, NA: 0.37, OD: 200 μm) which the other end connected to fiber photometry apparatus. After 30 min habituation, the voles were subjected to consolation test as described above (exposure to their shocked partners) and the behavioral recording consisted of allogrooming, sniffing, social approaching, self-grooming and running. During the test, if a specific behavior occurred no more than four times within 1 h, the record was replicated in the following day.

Fiber photometry signals were processed with custom-written MATLAB software. Briefly, all the data were segmented based on the behavioral events and baseline phase. The change in fluorescence (ΔF/F) was calculated as (F–F0)/F0, where F0 represents the baseline fluorescence signal averaged over a 10 s-long control time window. We first segmented the data based on the behavioral events. Then, we calculated the average 5-HT and calcium signals in both the pre- and post-phases (2/8 s). The time window was determined according to the duration of a behavior bout. The response elicited during a behavior was calculated as the average ΔF/F during all trials of a particular behavior. The peak response during a behavior was calculated as the maximum ΔF/F during the behavior minus F0.

For specificity verification of 5-HT sensors in Figure 5—figure supplement 2, a 5-HT receptor antagonist metergoline (Met) was injected (i.p., 4 mg/kg) 20 min before the fiber photometry experiments. The timing and dosing were chosen according to previous studies (Onasanwo et al., 2016; Wan, et al 2020).

### *In vitro* electrophysiological recordings

To verify the optogenetic and chemogenetic manipulations, we performed *in vitro* whole-cell patch-clamp recordings from DR neurons. Neurons expressing ChR2, eNpHR3.0, hM3Dq and hM4Di were visually identified by fluorescence of mCherry. The voles were anesthetized with isoflurane. Brains were quickly dissected and 300 μm coronal slices containing the DRN were prepared in a chamber filled with ACSF (32-34℃) using vibratome (Campden 7000 smz). The recordings were obtained using a Multiclamp 700B amplifier, filtered at 5 kHz and sampled at 10 kHz with Digidata 1440A. Clampex 10.5 was used for analysis.

Current-clamp recordings were performed to measure evoked action potentials. For photoactivation and photoinhibition, the light protocols used during behavioral tests were delivered through a 200 μm optical fiber close to recorded neurons. For CNO activation, spontaneous firing of action potentials in the cell was recorded in current clamp mode at −60 mV. After 5 min of recording, the slices were perfused with 10 μM CNO. The total recording time for each cell was 10 min. For CNO inhibition, we applied currents in steps of 25 pA, ranging from 0 pA to 250 pA. Neurons were allowed to recover for 10 min then perfused with 10 μM CNO. The same current procedure was performed. Afterwards, the CNO was removed by washes with ASCF and cells were recorded for another 10 min.

### Data analysis

All data are represented as the mean ± SE. All data were assessed for normality using a one-sample Kolmogorov-Smirnov test, and the Levene’s test was used to confirm homogeneity of variance. Comparisons between two groups were performed by unpaired or paired *t* tests. Comparisons among three or more groups of different animals were performed using one-way ANOVA. Two-way ANOVAs or two-way repeated measures ANOVAs were used to compare multiple groups under multiple testing conditions when appropriate. Post hoc comparisons were conducted using Tukey. To confirm the significance and calculate the effect size, Bayesian *t* test (paired or unpaired) and Bayesian ANOVA were conducted for all the comparisons using default priors. For more detailed information about the Bayesian test, please refer to Dr. Keysers’s work (Keysers et al., 2020). The data were analyzed using JASP 14.0 and are presented as the mean ± SE. The significant level was set at *P* < 0.05. The detailed analysis method in each figure and the statistical quantifications are presented in Supplementary 2.

## ACKNOWLEDGMENTS AND DISCLOSURES

This research was supported by the National Natural Science Foundation of China (31970424, 31670421, 31372213 and 31772473), the Natural Science Basic Research Plan in Shaanxin Province of China (2018JM3032) and the Fundamental Research Funds for Central University (GK201903059). We thank Yu-Ying Yang, Xin Zhang and Yi-Xin Feng for assistance in conducting experiments and caring for voles. The authors declare no conflict of interest.

## AUTHOR CONTRIBUTIONS

Prof. F.D.T. designed the study; L.F.L. conducted the majorities of experiments and wrote the original draft; L.Z.Z participated in the electrophysiological experiment; Z.X.H. and R.J. discussed the results and provided constructive comments; Y.T.Z participated in the fiber photometry studies. H.M., Y.F.X, W.Y., W.J.H., Z.J.L. and L.Y.T. participated in the behavior study and helped to collect and analyze the data. All authors contributed to and have approved the final manuscript.

## APPENDIX

**Supplementary 1. Supplemental figures.**

**Supplementary 2. Analysis methods and the statistical quantifications**

## SUPPLEMENTARY 1. SUPPLEMENTAL FIGURES.

**Figure 1—figure supplement 1.**
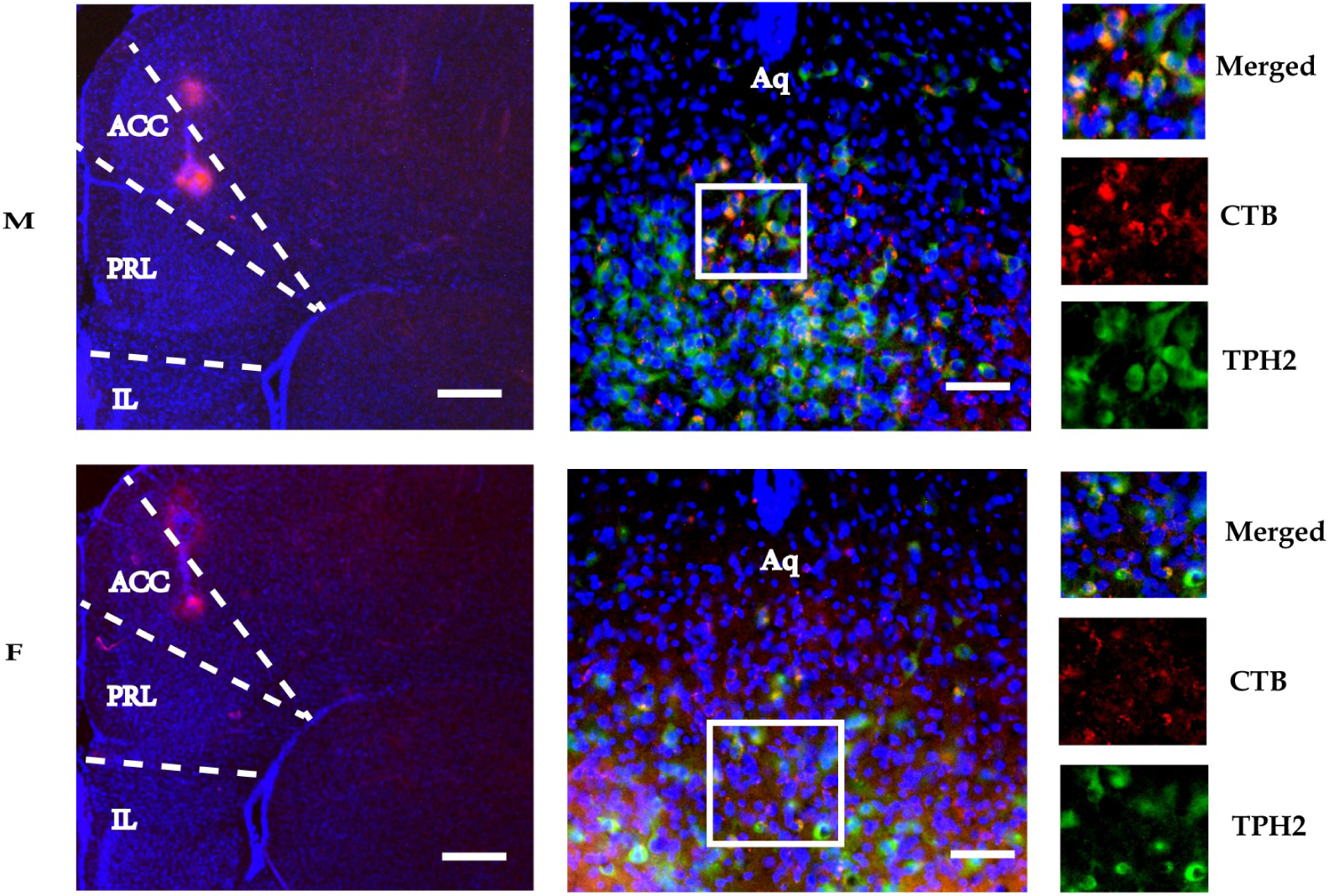
The histology of CTB injecting into the right ACC of male (up row) and female voles (down row). The right panels showed colocalization of TPH2^+^ neurons (green) and CTB (red) in the DR (× 200). M: male; F: female; ACC: anterior cingulate cortex; DR: dorsal raphe nucleus; TPH2: tryptophan hydroxylase 2; Aq: aqueduct.

**Figure 1—figure supplement 2.**
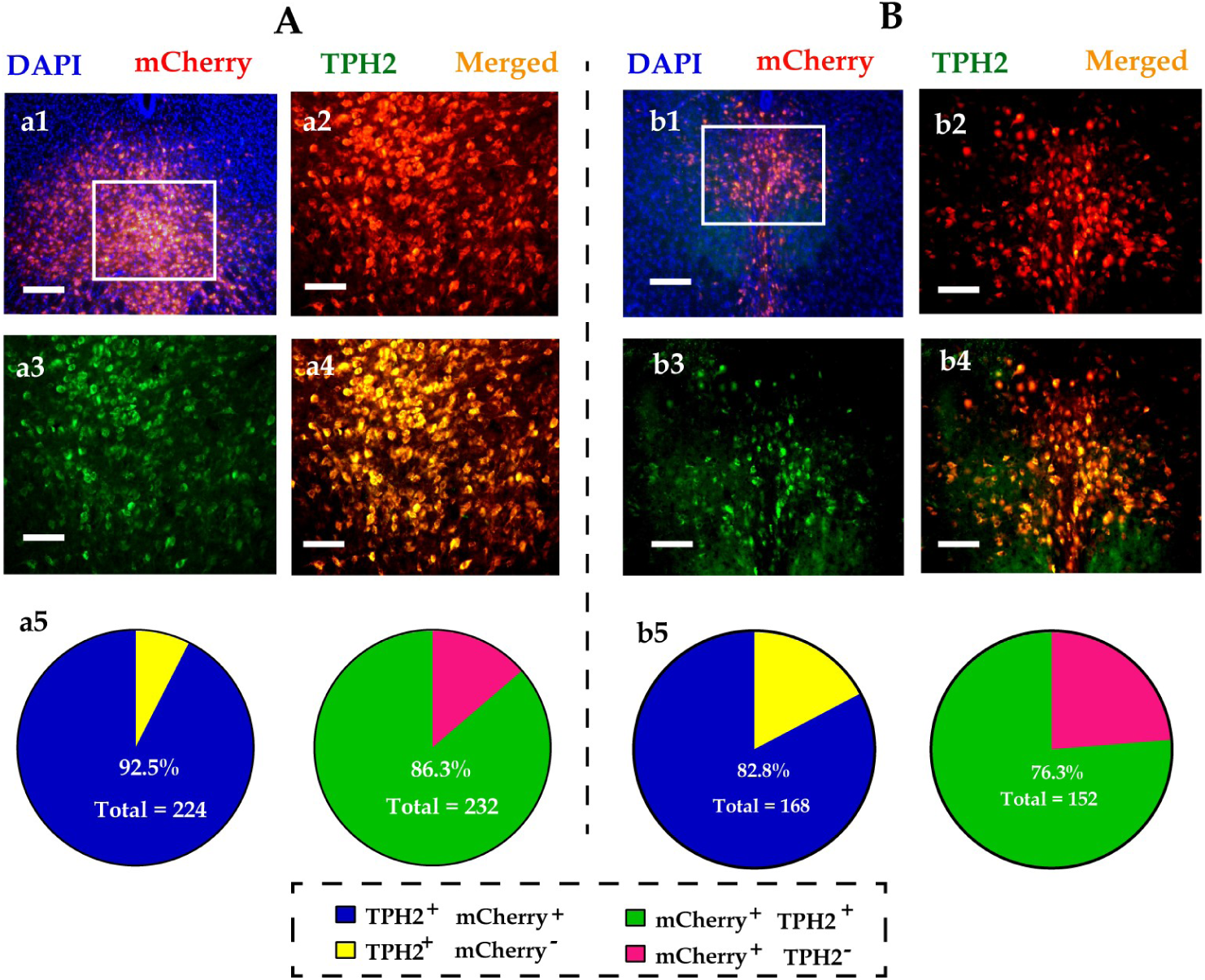
Immunohistological images showing colocalization of opsins (mCherry, red), TPH2+ neurons (green) and DAPI (blue) in the DR of males (A) and (B) females voles. (a1 & b1): merged image of DAPI and mCherry (× 100); (a2-a4, b2-b4): amplified images in the left box showing the mCherry, TPH2 and the colocaliztion of mCherry and TPH2 (× 200); (a5, b5): quantification rates of mCherry neurons colabled with TPH2 (left pies), and TPH2 neurons colabled with mCherry (right pies), *n* = 3 in each sex. DR: dorsal raphe nucleus; TPH2: tryptophan hydroxylase 2.

**Figure 1—figure supplement 3.**
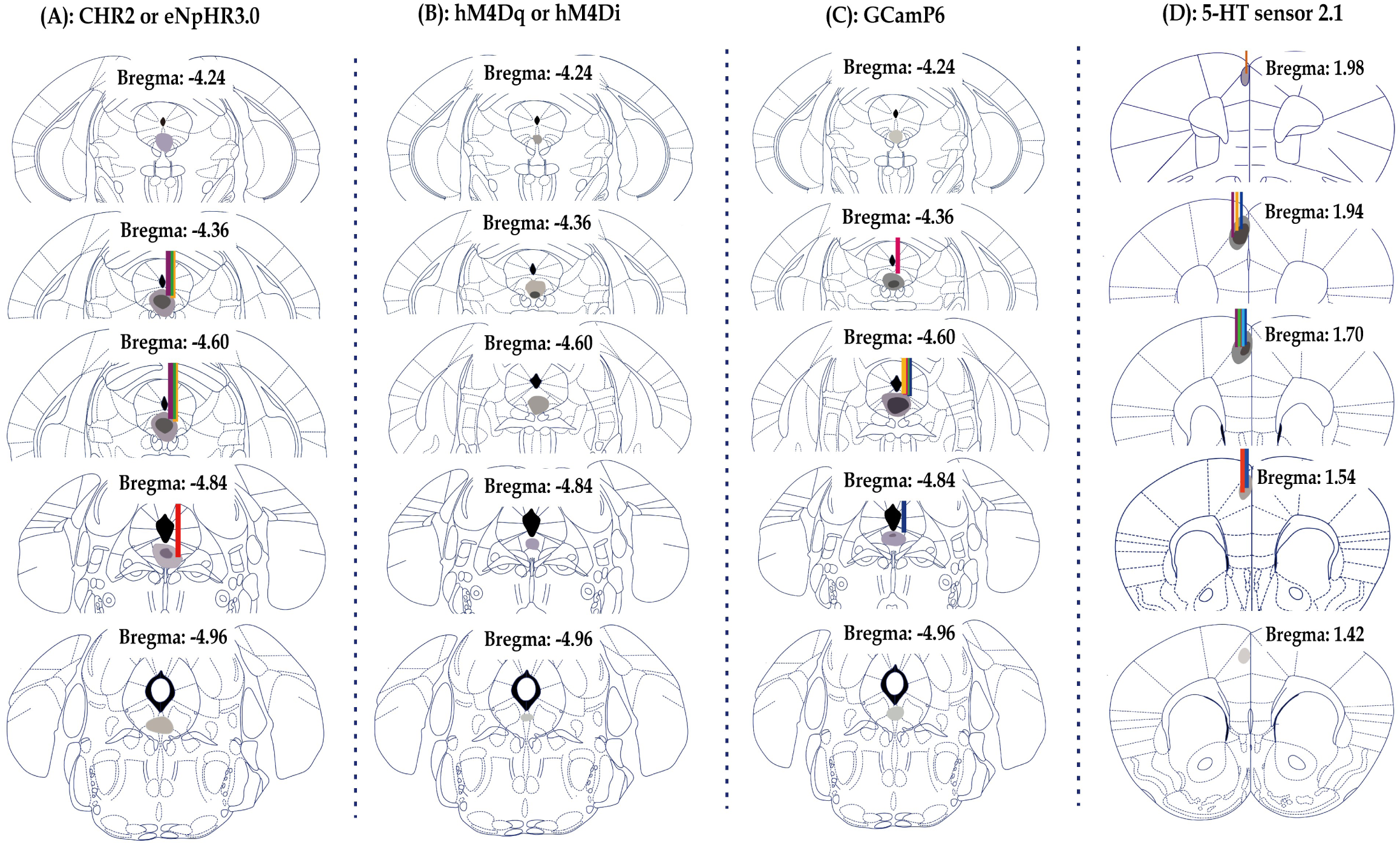
Schematics depict virus spread (shades) and optic fiber placements (lines) for recording and functional manipulation experiments, related to figures 1, 3, 4 and 5 (A, B C and D, respectively).

**Figure 1—figure supplement 4.**
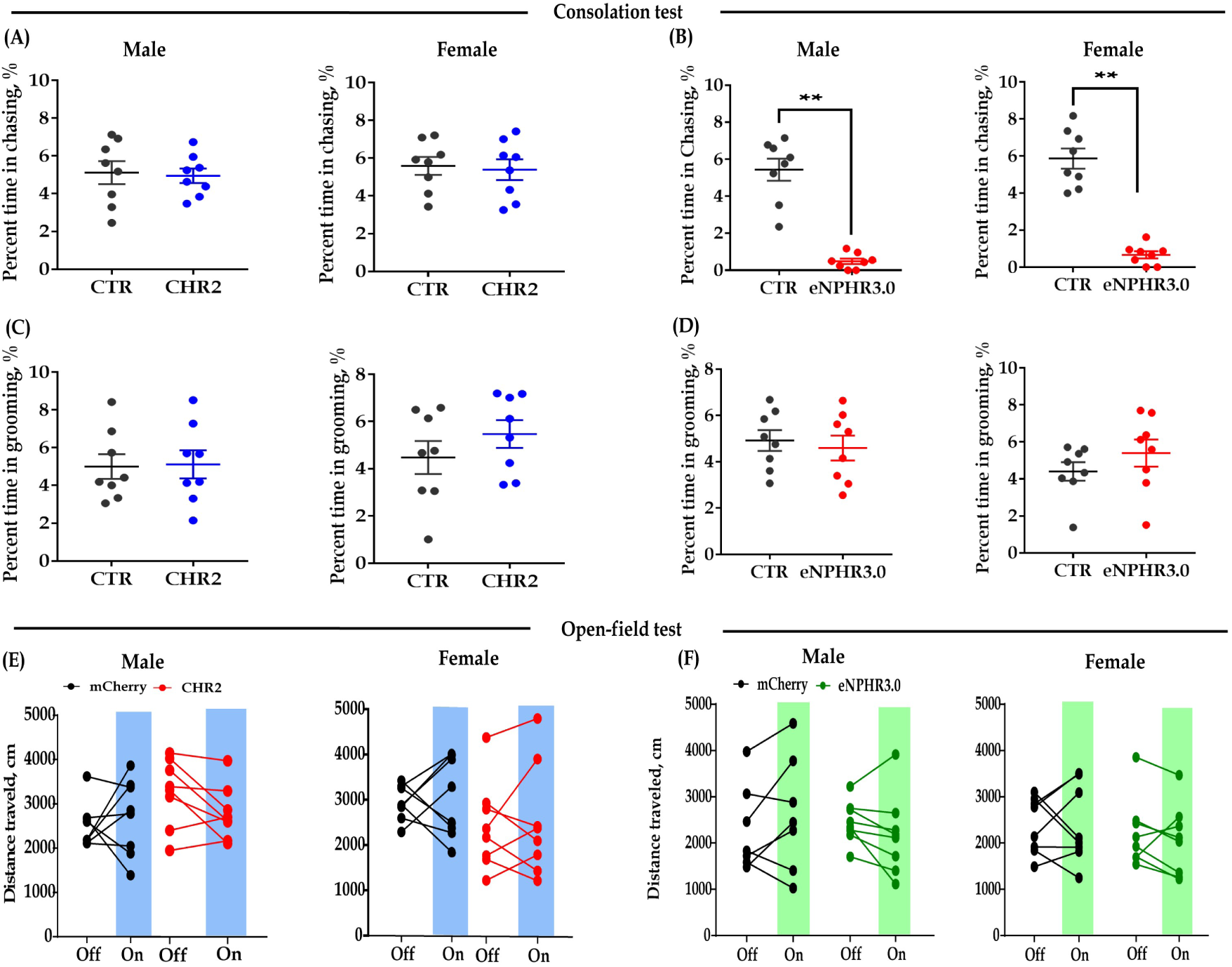
Effect of bidirectional optogenetic modulation of DR 5-HT neuron activities in the DR-ACC neural circuit on some control behaviors. Quantification of chasing time (A & B) and grooming time (C & D) in the consolation test; and the distance traveled in the open-field test (E & F). Data are presented as mean ± SE, *n* = 7–8 in each group, \*\**P* < 0.01. For A-D, Independent samples *t* test and Bayesian samples t-test; for E-F, two-way repeated ANOVA and two-way Bayesian Repeated Measures ANOVA (light as within subject factors). ACC: anterior cingulate cortex; CTR: control; DR: dorsal raphe nucleus.

**Figure 1—figure supplement 5.**
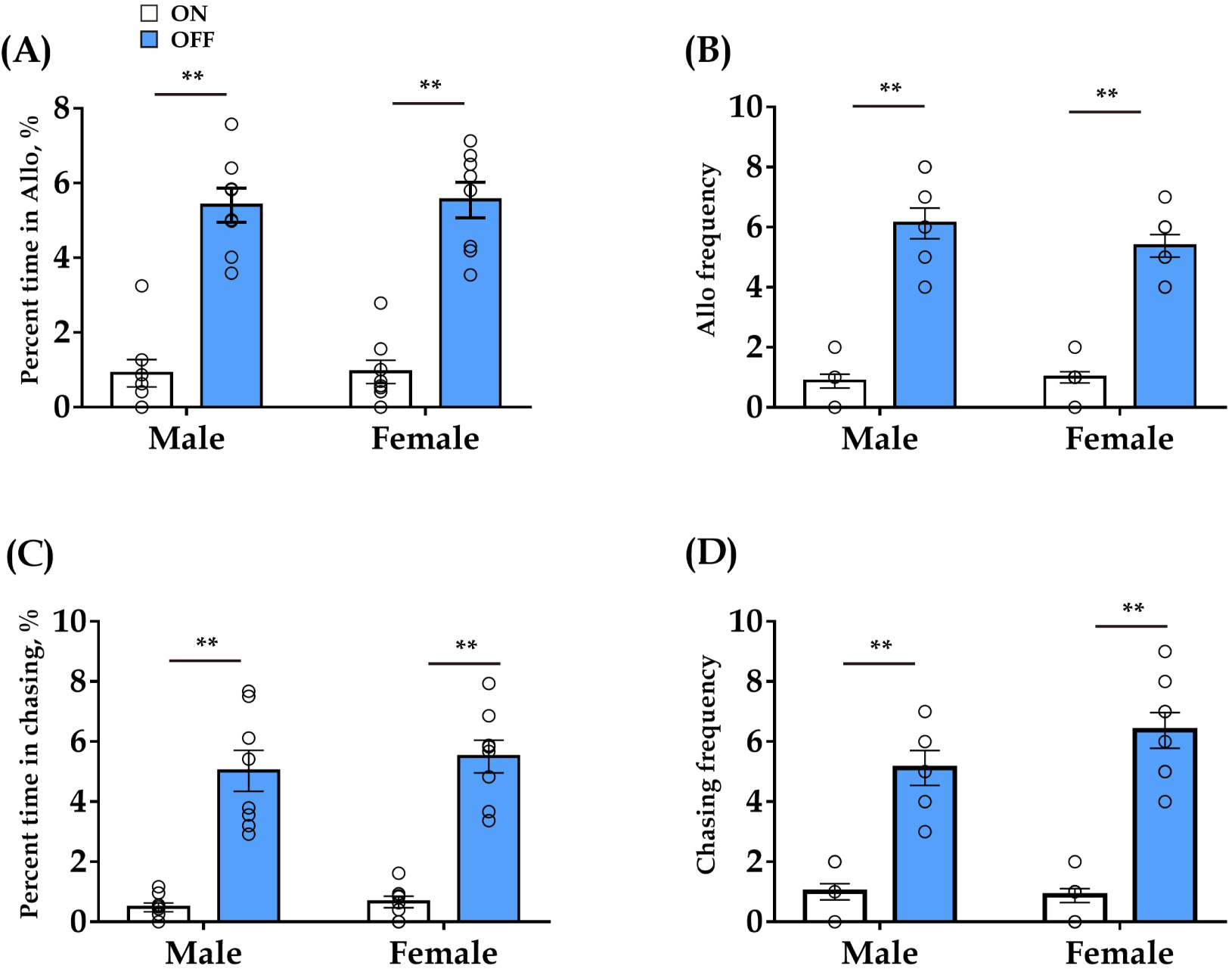
Effect of optogenetic inhibition of DR 5-HT neurons in the DR-ACC neural circuit does not elicit long-lasting effects (< 24 h) on allogrooming and chasing behavior in the consolation test. The data are derived from Figure 1H. (A): Time spent in allogrooming; (B) allogrooming frequency; (C): time spent in chasing and chasing frequency (D). Data are presented as mean ± SE, *n* = 8 in each group; Paired sample t-test and Bayesian Paired samples *t*-test; \*\**P* < 0.01. ACC: anterior cingulate cortex; DR: dorsal raphe nucleus; CTR: control.

**Figure 2—figure supplement 1.**
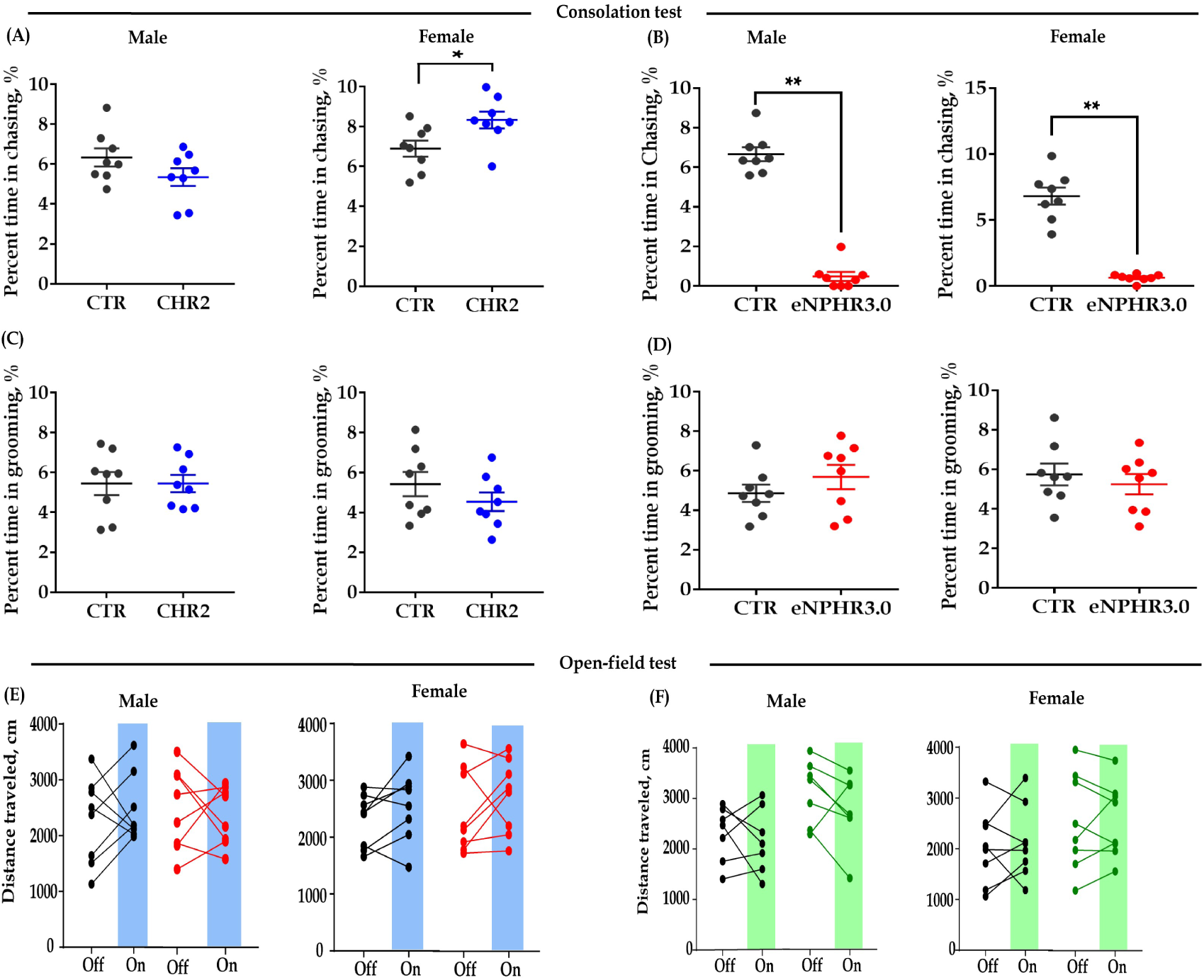
Effect of bidirectional optogenetic modulation of ACC 5-HT terminals in the DR-ACC neural circuit on some control behaviors. Quantification of chasing time (A & B) and grooming time (C & D) in the consolation test and the distance traveled in the open-field test (E & F). Data are presented as mean ± SE, *n* = 7–8 in each group, \*\**P* < 0.01. For A-D, Independent samples *t* test and Bayesian samples t-test; for E-F, two-way repeated ANOVA and two-way Bayesian Repeated Measures ANOVA (light as within subject factors). ACC: anterior cingulate cortex; DR: dorsal raphe nucleus; CTR: control.

**Figure 2—figure supplement 2.**
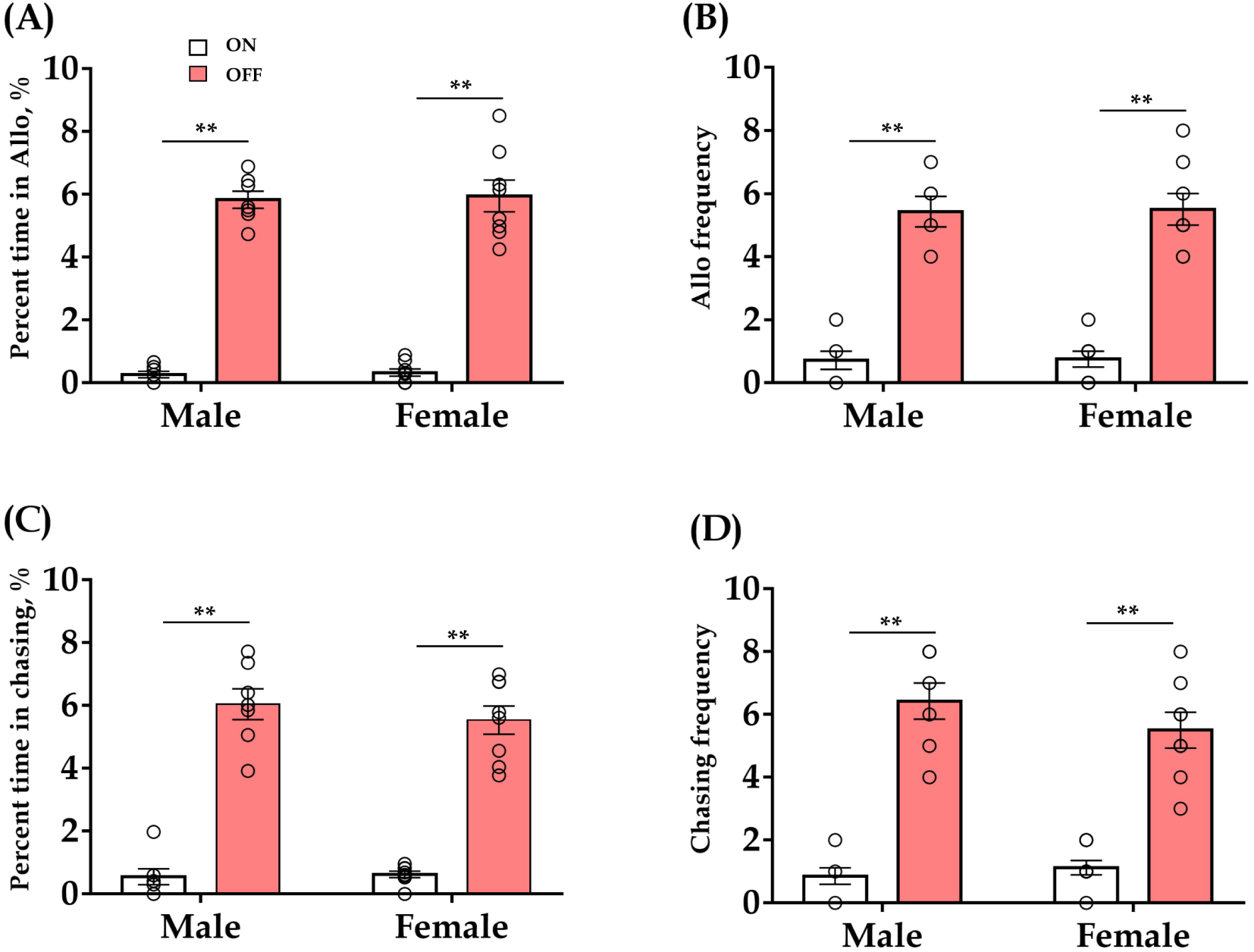
Effect of optogenetic inhibition of ACC 5-HT terminals in the DR-ACC neural circuit does not elicit long-lasting effects (< 24 h) on allogrooming and chasing behaviors in the consolation test. The data are derived from Figure 2D. (A): Time spent in allogrooming; (B) allogrooming frequency; (C): time spent in chasing; (D): chasing frequency. Data are presented as mean ± SE, *n* = 7–8 in each group; Paired sample t-test and Bayesian Paired samples t-test; \*\**P* < 0.01. ACC: anterior cingulate cortex; DR: dorsal raphe nucleus; CTR: control; Allo: allogrooming.

**Figure 3—figure supplement 1.**
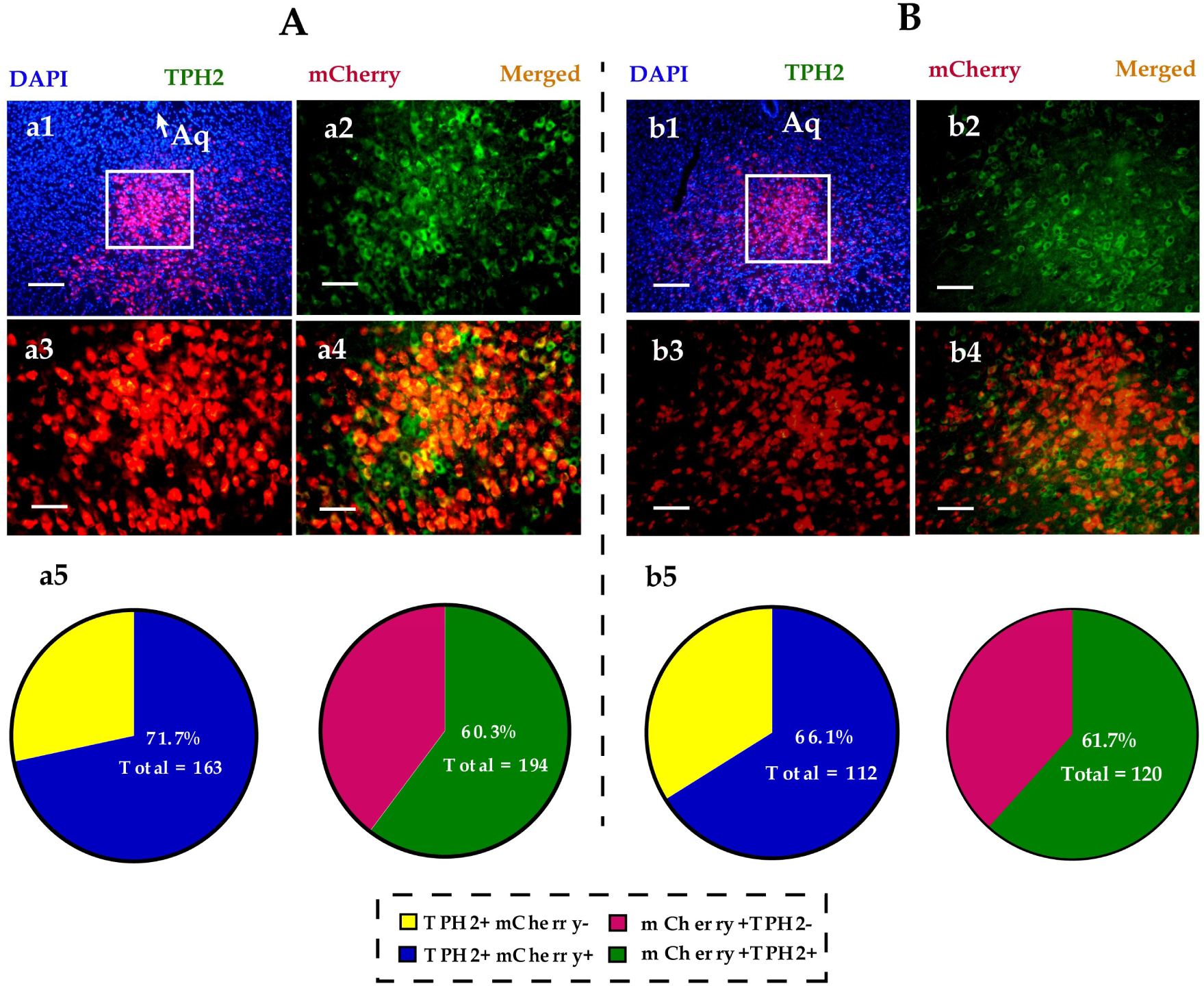
Immunohistological image showing colocalization of DREADD (mCherry, red), TPH2+ neurons (green) and DAPI (blue) in the DR of male (A) and (B) female voles. (a1 & b1): merged image of DAPI and mCherry (× 100); (a2-a4, b2-b4): amplified images in the left box showing the mCherry, TPH2 and the colocaliztion of mCherry and TPH2 (× 200); (a5, b5): quantification rates of mCherry neurons colabled with TPH2 (left pies), and TPH2 neurons colabled with mCherry (right pies), *n* = 3 in each sex. TPH2: tryptophan hydroxylase 2.

**Figure 3—figure supplement 2.**
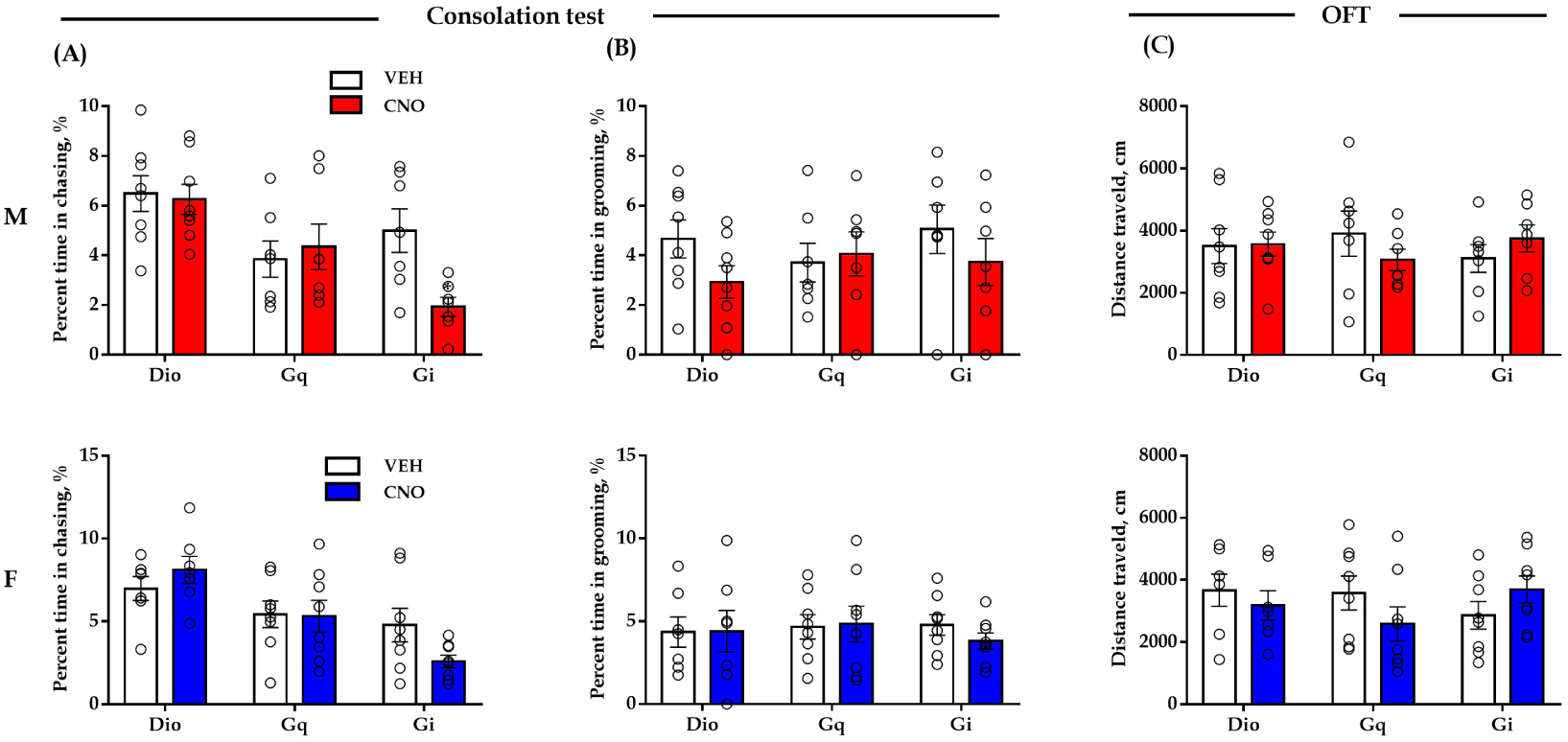
Effect of chemogenetic modulation of DR 5-HT neuron activities in the DR-ACC neural circuit on some control behaviors. Quantification of chasing time (A) and grooming time (B) in the consolation test, and distance traveled in the open-field test (C). Data are presented as mean ± SE, *n* = 7–8 in each group; two-way repeated measures ANOVA along with two-way Bayesian repeated measures ANOVA. ACC: anterior cingulate cortex; DR: dorsal raphe nucleus; M: male; F: female; OFT: open-field test.

**Figure 4—figure supplement 1.**
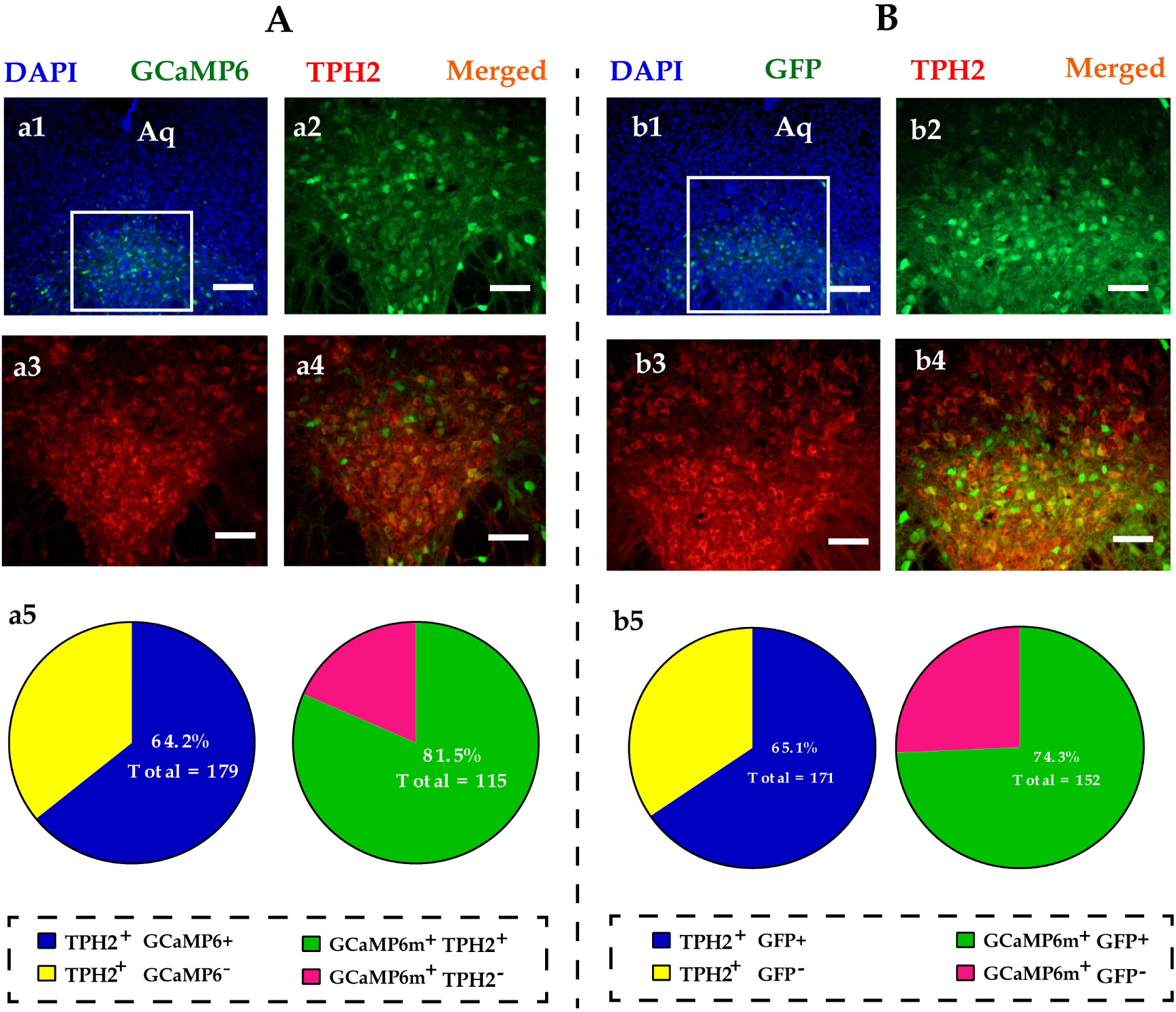
Representative viral infection images of GCaMp6. (a1): Immunohistological image showing GCaMP6 expression in the DR (× 100); (a2-a4): amplified images in the left box showing the GCaMP6 (green), TPH2 (red) and the colocaliztion (yellow) of the two (× 200); (a5): quantification rates of GCaMP6 neurons colabled with TPH2 (left pie), and TPH2 neurons colabled with GCaMP6 (right pie), *n* = 3; (b1): immunohistological image showing GFP expression in the DR (× 100); (b2-b4): amplified images in the left box showing the GFP (green), TPH2 (red) and the colocaliztion (yellow) of the two (× 200); (b5): quantification rates of GFP neurons colabled with TPH2 (left pie), and TPH2 neurons colabled with GFP (right pie), *n* = 3.

**Figure 4—figure supplement 2.**
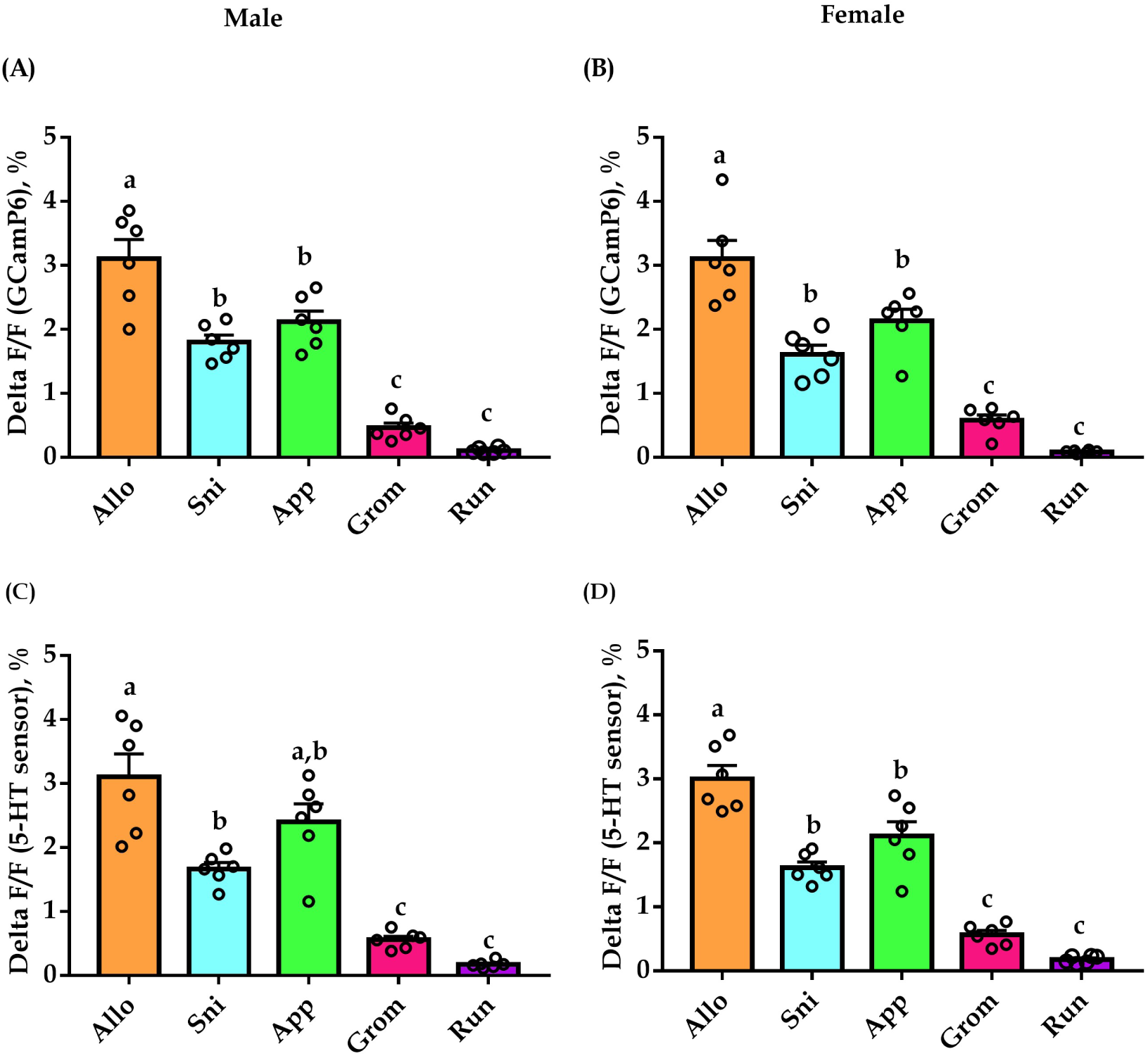
Peak GCaMP6 (A & B) and 5-HT sensor (C & D) fluorescence values during a few of behaviors in both male and female voles. Data are presented as mean ± SE, *n* = 6 in each group. Groups not sharing the same letter significantly differ from each other; one-way ANOVA along with one-way Bayesian ANOVA. Allo: allogrooming; Sni: sniffing; App: social approaching; Gro: selfgrooming; Run: running.

**Figure 4—figure supplement 3.**
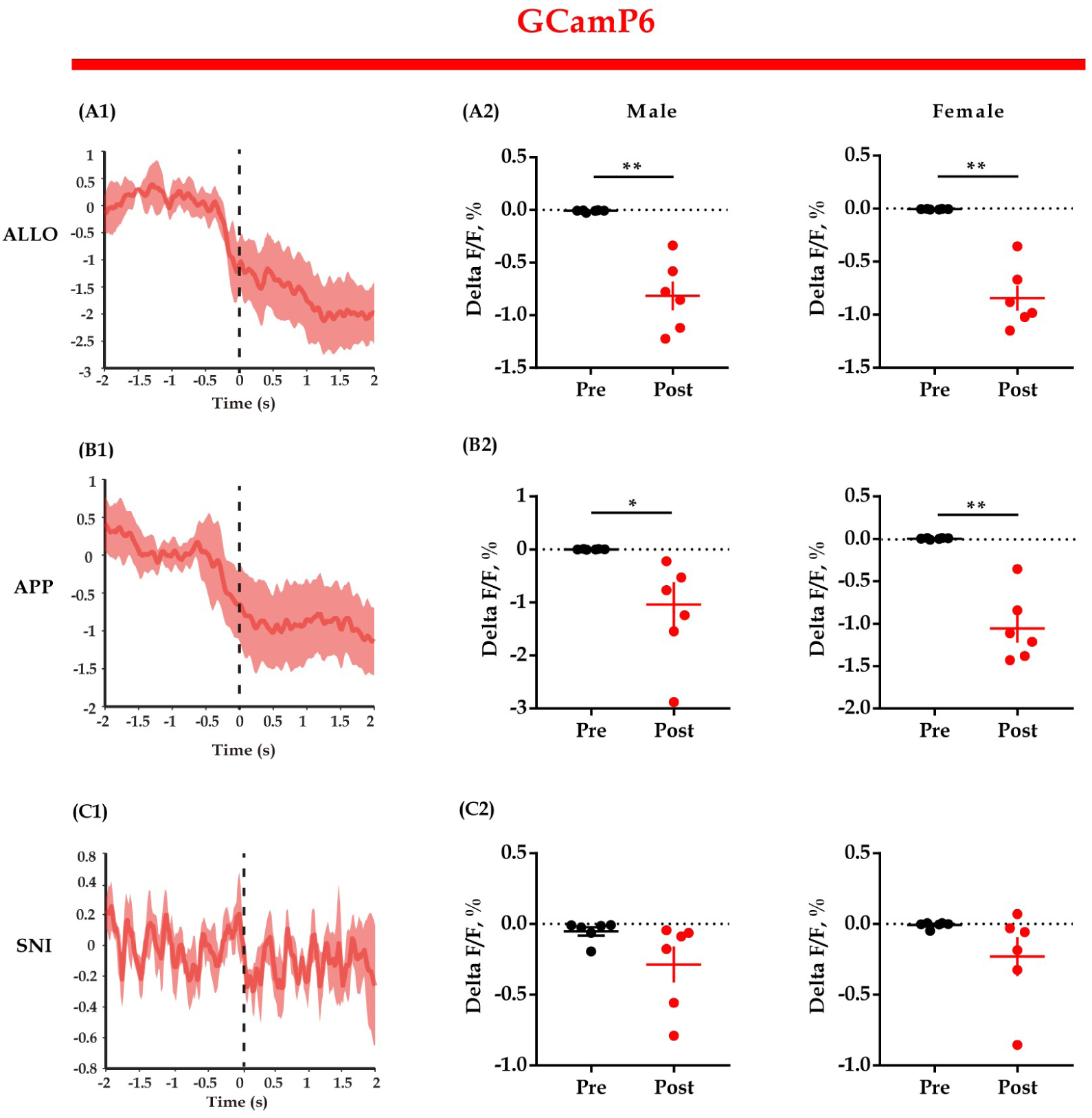
GCaMP6 fluorescent signals align to the end of some behaviors. (A1, B1, C1): Representative peri-event plot of GCaMP6 fluorescence signals aligned to the end of allogrooming, social approaching and sniffing (for all peri-event plots, the red line denotes the mean signals of 4-6 bouts of behaviors, whereas the red shaded region denotes the SE); (A2, B2, C2): quantification of change in GCaMP6 fluorescence signals. Data are presented as mean ± SE, \**P* < 0.05, \*\**P* < 0.01, *n* = 6 in each group; paired samples *t* test along with Bayesian paired samples *t*-Test. ALLO: allogrooming; APP: approaching; SNI: sniffing.

**Figure 5—figure supplement 1.**
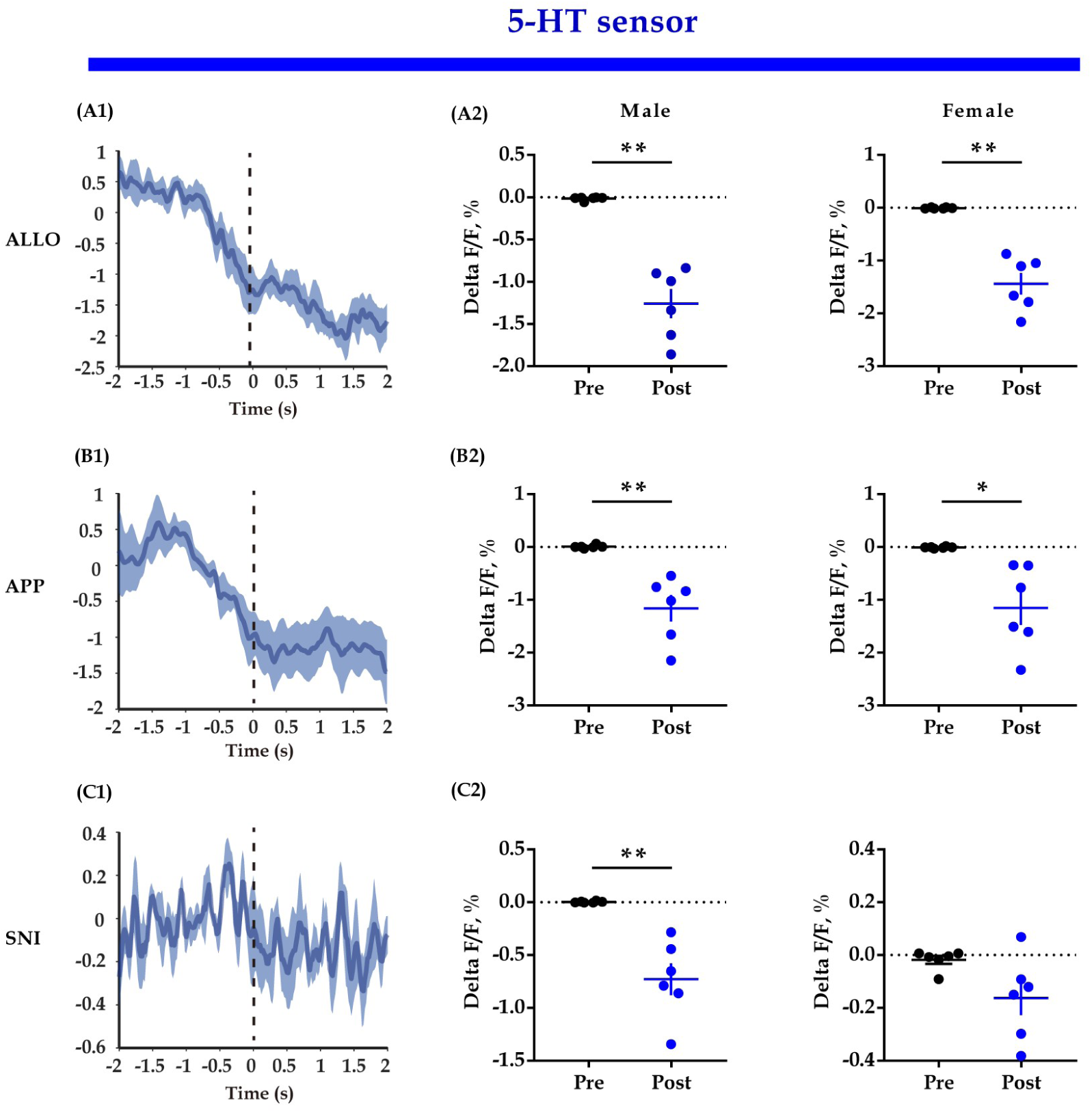
5-HT sensor fluorescent signals align to the end of some behaviors. (A1, B1, C1): Representative peri-event plot of 5-HT sensor fluorescence signals aligned to the end of allogrooming, social approaching and sniffing (for all peri-event plots, the blue line denotes the mean signals of 4-6 bouts of behaviors, whereas the blue shaded region denotes the SE); (A2, B2, C2): quantification of change in 5-HT sensor fluorescence signals. Data are presented as mean ± SE, \**P* < 0.05, \*\**P* < 0.01, *n* = 6 in each group; simple Paired *t* test along with Bayesian Paired Samples *t*-Test. ALLO: allogrooming; APP: approaching; SNI: sniffing.

**Figure 5—figure supplement 2.**
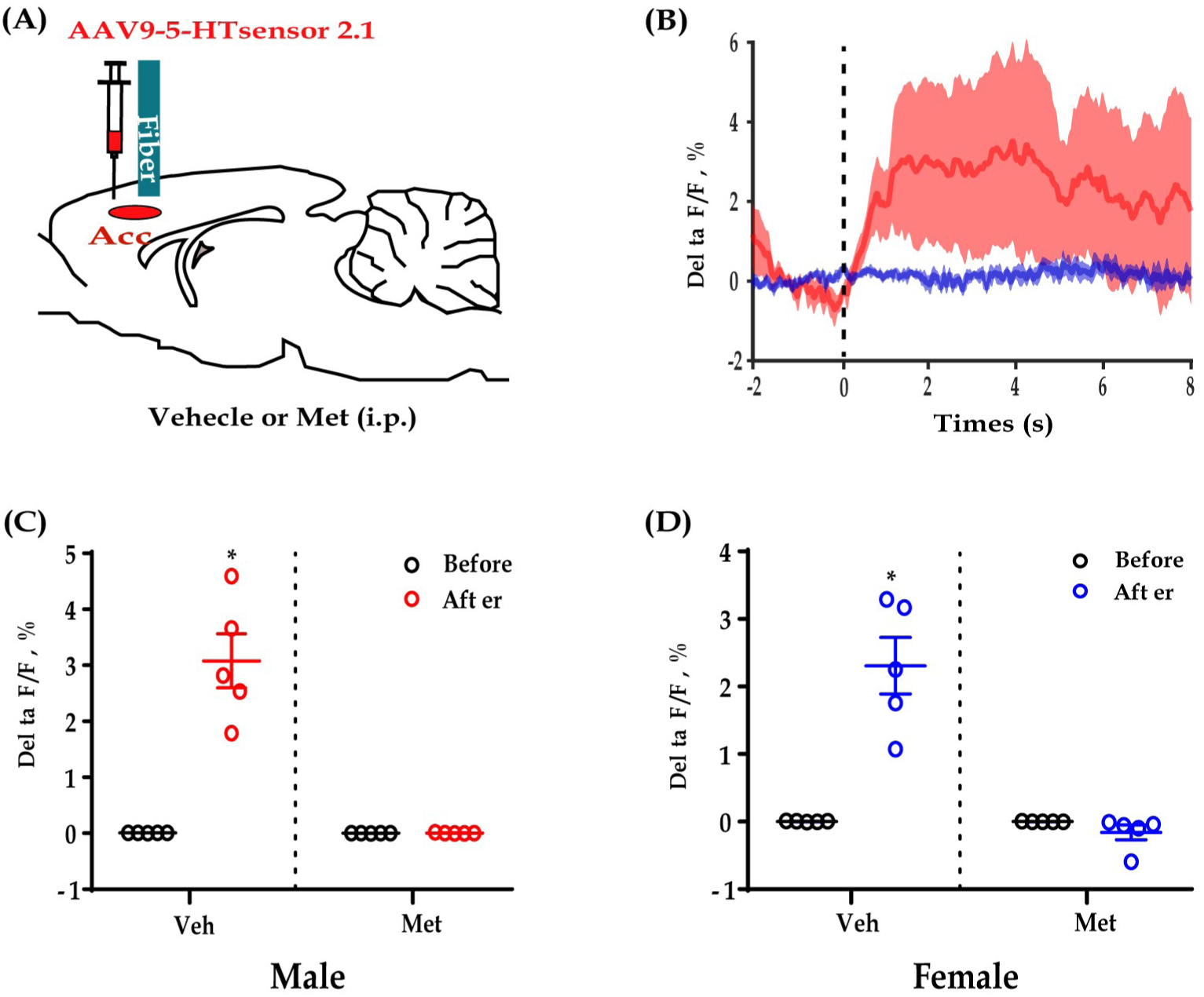
5-HT sensor fluoresce changes during allogrooming could be blocked by Met. (A): Schematic diagrams; (B): combined individual 5-HT sensor fluorescence traces during allogrooming in response to Met (blue) or Veh (red); (C, D): quantification of 5-HT sensor fluorescence signals during allogrooming in response to Met or Veh in male (C) and female subjects. Data are presented as mean ± SE, \*\**P* < 0.01, *n* = 6 in each group; two-way Repeated measures ANOVA and two-way Bayesian repeated measures ANOVA. Met: metergoline, a 5-HT receptor antagonist; Veh: vehicle.

**Figure 6—figure supplement 1.**
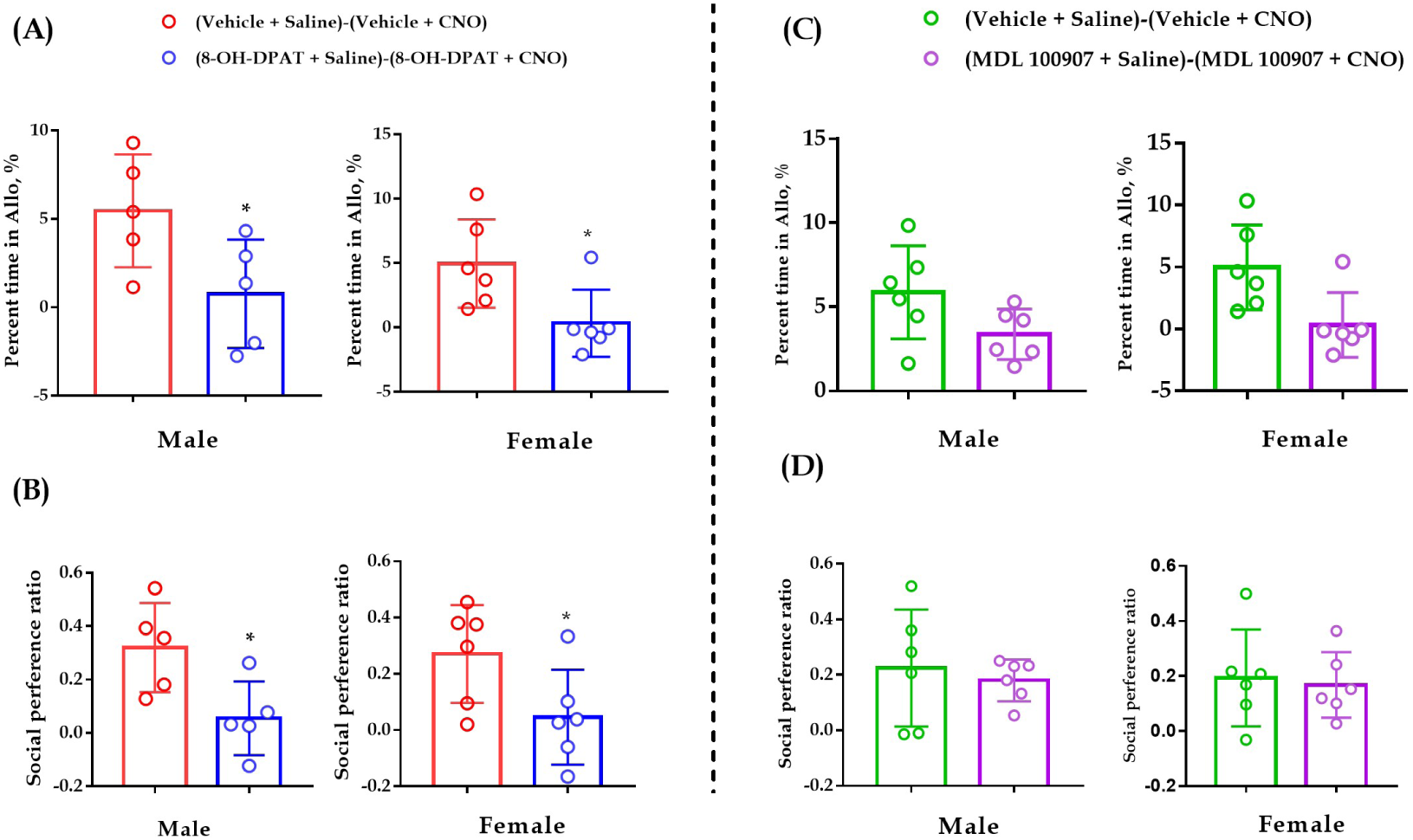
Comparisons of ‘Vehicle+Saline - Vehicle+CNO’ vs ‘8-OH-DPAT+Saline - 8-OH-DPAT+CNO’ (A & B) and ‘Vehicle+Saline - Vehicle+CNO’ vs ‘MDL 100907+Saline - MDL 100907+CNO’ (C & D). Independent samples *t*-test and Bayesian independent samples *t*-test*. N* = 5-6 in each group, \**P* < 0.05.

**Figure 6—figure supplement 2.**
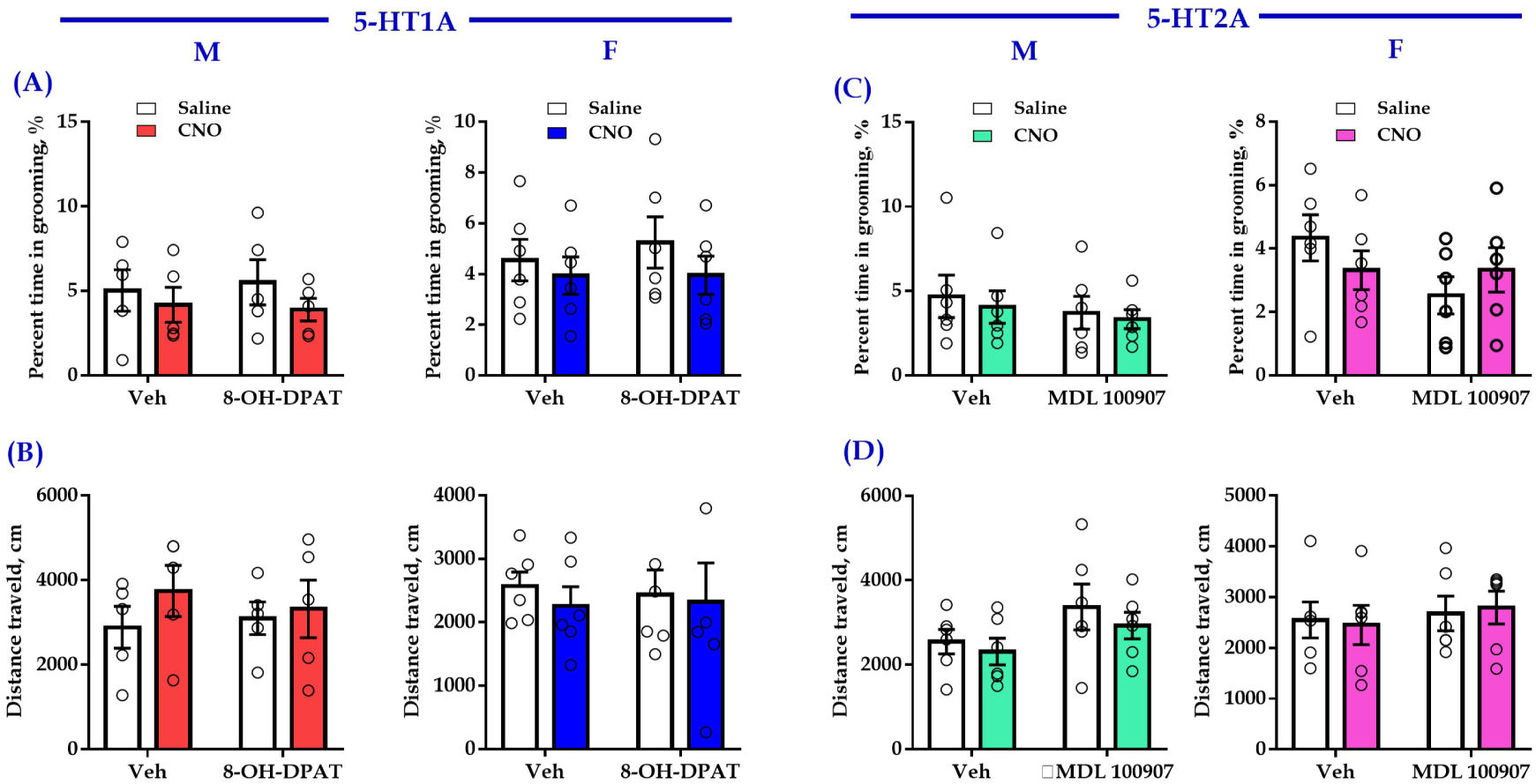
Chemogenetic inhibition of DR 5-HT neuron in the DR-ACC neural circuit along with intra-ACC injection of 5-HT1AR agonist (8-OH-DPAT) or 5-HT2AR antagonist (MDL 100907) had no significant effect on control behaviors of selfgrooming (A, C) and distance traveled in the open-field test (B, D). Two-way ANOVA and two-way Bayesian ANOVA*. N* = 5-6 in each group. ACC: anterior cingulate cortex; DR: dorsal raphe nucleus; M: male; F: female.

## REFERENCE

Arakawa, H., 2020. Somatosensorimotor and Odor Modification, Along with Serotonergic Processes Underlying the Social Deficits in BTBR T+ Itpr3(tf)/J and BALB/cJ Mouse Models of Autism. Neuroscience 445, 144–162.

Artigas, F., 2013. Serotonin receptors involved in antidepressant effects. Pharmacology & therapeutics 137, 119–131.

Bozzi, Y., Provenzano, G., Casarosa, S., 2018. Neurobiological bases of autism-epilepsy comorbidity: a focus on excitation/inhibition imbalance. The European journal of neuroscience 47, 534–548.

Burkett, J.P., Andari, E., Johnson, Z.V., Curry, D.C., de Waal, F.B., Young, L.J., 2016. Oxytocin-dependent consolation behavior in rodents. Science (New York, N.Y.) 351, 375–378.

Carhart-Harris, R.L., Nutt, D.J., 2017. Serotonin and brain function: a tale of two receptors. Journal of psychopharmacology (Oxford, England) 31, 1091–1120.

Carlyle, M., Stevens, T., Fawaz, L., Marsh, B., Kosmider, S., Morgan, C.J., 2019. Greater empathy in MDMA users. Journal of psychopharmacology (Oxford, England) 33, 295–304.

Celada, P., Puig, M.V., Artigas, F., 2013. Serotonin modulation of cortical neurons and networks. Frontiers in integrative neuroscience 7, 25.

Charnay, Y., Leger, L., 2010. Brain serotonergic circuitries. Dialogues in clinical neuroscience 12, 471–487.

Cooper, M.A., McIntyre, K.E., Huhman, K.L., 2008. Activation of 5-HT1A autoreceptors in the dorsal raphe nucleus reduces the behavioral consequences of social defeat. Psychoneuroendocrinology 33, 1236–1247.

Correia, P.A., Lottem, E., Banerjee, D., Machado, A.S., Carey, M.R., Mainen, Z.F., 2017. Transient inhibition and long-term facilitation of locomotion by phasic optogenetic activation of serotonin neurons. eLife 6.

de Waal, F.B., 2008. Putting the altruism back into altruism: the evolution of empathy. Annual review of psychology 59, 279–300.

de Waal, F.B.M., Preston, S.D., 2017. Mammalian empathy: behavioural manifestations and neural basis. Nature reviews. Neuroscience 18, 498–509.

Dolder, P.C., Grunblatt, E., Muller, F., Borgwardt, S.J., Liechti, M.E., 2017. A Single Dose of LSD Does Not Alter Gene Expression of the Serotonin 2A Receptor Gene (HTR2A) or Early Growth Response Genes (EGR1-3) in Healthy Subjects. Frontiers in pharmacology 8, 423.

Dolen, G., Darvishzadeh, A., Huang, K.W., Malenka, R.C., 2013. Social reward requires coordinated activity of nucleus accumbens oxytocin and serotonin. Nature 501, 179–184.

Faye, C., Hen, R., Guiard, B.P., Denny, C.A., Gardier, A.M., Mendez-David, I., David, D.J., 2020. Rapid Anxiolytic Effects of RS67333, a Serotonin Type 4 Receptor Agonist, and Diazepam, a Benzodiazepine, Are Mediated by Projections From the Prefrontal Cortex to the Dorsal Raphe Nucleus. Biological psychiatry 87, 514–525.

Feng, J., Zhang, C., Lischinsky, J.E., Jing, M., Zhou, J., Wang, H., Zhang, Y., Dong, A., Wu, Z., Wu, H., Chen, W., Zhang, P., Zou, J., Hires, S.A., Zhu, J.J., Cui, G., Lin, D., Du, J., Li, Y., 2019. A Genetically Encoded Fluorescent Sensor for Rapid and Specific In Vivo Detection of Norepinephrine. Neuron 102, 745–761 e748.

Field, T., Diego, M., Hernandez-Reif, M., 2009. Depressed mothers’ infants are less responsive to faces and voices. Infant behavior & development 32, 239–244.

Flanigan, M.E., Aleyasin, H., Li, L., Burnett, C.J., Chan, K.L., LeClair, K.B., Lucas, E.K., Matikainen-Ankney, B., Durand-de Cuttoli, R., Takahashi, A., Menard, C., Pfau, M.L., Golden, S.A., Bouchard, S., Calipari, E.S., Nestler, E.J., DiLeone, R.J., Yamanaka, A., Huntley, G.W., Clem, R.L., Russo, S.J., 2020. Orexin signaling in GABAergic lateral habenula neurons modulates aggressive behavior in male mice. Nature neuroscience.

Fu, W., Le Maitre, E., Fabre, V., Bernard, J.F., David Xu, Z.Q., Hokfelt, T., 2010. Chemical neuroanatomy of the dorsal raphe nucleus and adjacent structures of the mouse brain. The Journal of comparative neurology 518, 3464–3494.

Fukumoto, K., Fogaça, M.V., Liu, R.J., Duman, C.H., Li, X.Y., Chaki, S., Duman, R.S., 2020. Medial PFC AMPA receptor and BDNF signaling are required for the rapid and sustained antidepressant-like effects of 5-HT(1A) receptor stimulation. Neuropsychopharmacology : official publication of the American College of Neuropsychopharmacology 45, 1725–1734.

Fukumoto, K., Iijima, M., Chaki, S., 2014. Serotonin-1A receptor stimulation mediates effects of a metabotropic glutamate 2/3 receptor antagonist, 2S-2-amino-2-(1S,2S-2-carboxycycloprop-1-yl)-3-(xanth-9-yl)propanoic acid (LY341495), and an N-methyl-D-aspartate receptor antagonist, ketamine, in the novelty-suppressed feeding test. Psychopharmacology 231, 2291–2298.

Garcia-Garcia, A.L., Canetta, S., Stujenske, J.M., Burghardt, N.S., Ansorge, M.S., Dranovsky, A., Leonardo, E.D., 2018. Serotonin inputs to the dorsal BNST modulate anxiety in a 5-HT1A receptor-dependent manner. Molecular psychiatry 23, 1990–1997.

Gong, P., Liu, J., Blue, P.R., Li, S., Zhou, X., 2015. Serotonin receptor gene (HTR2A) T102C polymorphism modulates individuals’ perspective taking ability and autistic-like traits. Frontiers in human neuroscience 9, 575.

Gyurak, A., Haase, C.M., Sze, J., Goodkind, M.S., Coppola, G., Lane, J., Miller, B.L., Levenson, R.W., 2013. The effect of the serotonin transporter polymorphism (5-HTTLPR) on empathic and self-conscious emotional reactivity. Emotion (Washington, D.C.) 13, 25–35.

Heifets, B.D., Malenka, R.C., 2016. MDMA as a Probe and Treatment for Social Behaviors. Cell 166, 269–272.

Horie, K., Inoue, K., Suzuki, S., Adachi, S., Yada, S., Hirayama, T., Hidema, S., Young, L.J., Nishimori, K., 2019. Oxytocin receptor knockout prairie voles generated by CRISPR/Cas9 editing show reduced preference for social novelty and exaggerated repetitive behaviors. Hormones and behavior 111, 60–69.

Ishii, H., Ohara, S., Tobler, P.N., Tsutsui, K., Iijima, T., 2015. Dopaminergic and serotonergic modulation of anterior insular and orbitofrontal cortex function in risky decision making. Neuroscience research 92, 53–61.

Keysers, C., Gazzola, V., Wagenmakers, E.J., 2020. Using Bayes factor hypothesis testing in neuroscience to establish evidence of absence. Nature neuroscience 23, 788–799.

Kim, B.S., Lee, J., Bang, M., Seo, B.A., Khalid, A., Jung, M.W., Jeon, D., 2014. Differential regulation of observational fear and neural oscillations by serotonin and dopamine in the mouse anterior cingulate cortex. Psychopharmacology 231, 4371–4381.

Knapska, E., Mikosz, M., Werka, T., Maren, S., 2010. Social modulation of learning in rats. Learning & memory (Cold Spring Harbor, N.Y.) 17, 35–42.

Li, L.F., Yuan, W., He, Z.X., Ma, H., Xun, Y.F., Meng, L.R., Zhu, S.J., Wang, L.M., Zhang, J., Cai, W.Q., Zhang, X.N., Guo, Q.Q., Lian, Z.M., Jia, R., Tai, F.D., 2020. Reduced Consolation Behaviors in Physically Stressed Mandarin Voles: Involvement of Oxytocin, Dopamine D2, and Serotonin 1A Receptors Within the Anterior Cingulate Cortex. The international journal of neuropsychopharmacology 23, 511-523.

Li, L.F., Yuan, W., He, Z.X., Wang, L.M., Jing, X.Y., Zhang, J., Yang, Y., Guo, Q.Q., Zhang, X.N., Cai, W.Q., Hou, W.J., Jia, R., Tai, F.D., 2019. Involvement of oxytocin and GABA in consolation behavior elicited by socially defeated individuals in mandarin voles. Psychoneuroendocrinology 103, 14–24.

Li, Y., Zhong, W., Wang, D., Feng, Q., Liu, Z., Zhou, J., Jia, C., Hu, F., Zeng, J., Guo, Q., Fu, L., Luo, M., 2016. Serotonin neurons in the dorsal raphe nucleus encode reward signals. Nature communications 7, 10503.

Luo, M., Zhou, J., Liu, Z., 2015. Reward processing by the dorsal raphe nucleus: 5-HT and beyond. Learning & memory (Cold Spring Harbor, N.Y.) 22, 452–460.

Matsunaga, M., Ishii, K., Ohtsubo, Y., Noguchi, Y., Ochi, M., Yamasue, H., 2017. Association between salivary serotonin and the social sharing of happiness. PloS one 12, e0180391.

Meneses, A., Liy-Salmeron, G., 2012. Serotonin and emotion, learning and memory. Reviews in the neurosciences 23, 543–553.

Ohmura, Y., Tanaka, K.F., Tsunematsu, T., Yamanaka, A., Yoshioka, M., 2014. Optogenetic activation of serotonergic neurons enhances anxiety-like behaviour in mice. The international journal of neuropsychopharmacology 17, 1777–1783.

Onasanwo, S.A., Faborode, S.O., Ilenre, K.O., 2016. Antidepressant-like Potentials of Buchholzia Coriacea Seed Extract: Involvement of Monoaminergic and Cholinergic Systems, and Neuronal Density in the Hippocampus of Adult Mice. Nigerian journal of physiological sciences : official publication of the Physiological Society of Nigeria 31, 93–99.

Perez-Manrique, A., Gomila, A., 2018. The comparative study of empathy: sympathetic concern and empathic perspective-taking in non-human animals. Biological reviews of the Cambridge Philosophical Society 93, 248–269.

Pockros, L.A., Pentkowski, N.S., Swinford, S.E., Neisewander, J.L., 2011. Blockade of 5-HT2A receptors in the medial prefrontal cortex attenuates reinstatement of cue-elicited cocaine-seeking behavior in rats. Psychopharmacology 213, 307–320.

Puig, M.V., Artigas, F., Celada, P., 2005. Modulation of the activity of pyramidal neurons in rat prefrontal cortex by raphe stimulation in vivo: involvement of serotonin and GABA. Cerebral cortex (New York, N.Y. : 1991) 15, 1–14.

Puig, M.V., Gulledge, A.T., 2011. Serotonin and prefrontal cortex function: neurons, networks, and circuits. Molecular neurobiology 44, 449–464.

Santana, N., Artigas, F., 2017. Laminar and Cellular Distribution of Monoamine Receptors in Rat Medial Prefrontal Cortex. Frontiers in neuroanatomy 11, 87.

Santana, N., Bortolozzi, A., Serrats, J., Mengod, G., Artigas, F., 2004. Expression of serotonin1A and serotonin2A receptors in pyramidal and GABAergic neurons of the rat prefrontal cortex. Cerebral cortex (New York, N.Y. : 1991) 14, 1100–1109.

Tian, Z., Yamanaka, M., Bernabucci, M., Zhao, M.G., Zhuo, M., 2017. Characterization of serotonin-induced inhibition of excitatory synaptic transmission in the anterior cingulate cortex. Molecular brain 10, 21.

Vattikuti, S., Chow, C.C., 2010. A computational model for cerebral cortical dysfunction in autism spectrum disorders. Biological psychiatry 67, 672–678.

Walsh, J.J., Christoffel, D.J., Heifets, B.D., Ben-Dor, G.A., Selimbeyoglu, A., Hung, L.W., Deisseroth, K., Malenka, R.C., 2018. 5-HT release in nucleus accumbens rescues social deficits in mouse autism model. Nature 560, 589–594.

Wan JX, Peng WL, Li XL, Qian TG, Song K, Zeng JZ, et al. A genetically encoded GRAB sensor for measuring serotonin dynamics in vivo. Biorxiv. 2020; 38:569–580; posted February 25, 2020; doi: 10.1101/2020.02.24.962282.

Wang, L., Zhu, Z., Hou, W., Zhang, X., He, Z., Yuan, W., Yang, Y., Zhang, S., Jia, R., Tai, F., 2019. Serotonin Signaling Trough Prelimbic 5-HT1A Receptors Modulates CSDS-Induced Behavioral Changes in Adult Female Voles. The international journal of neuropsychopharmacology 22, 208–220.

Weber, E.T., Andrade, R., 2010. Htr2a Gene and 5-HT(2A) Receptor Expression in the Cerebral Cortex Studied Using Genetically Modified Mice. Frontiers in neuroscience 4.

Yagishita, S., 2020. Transient and sustained effects of dopamine and serotonin signaling in motivation-related behavior. Psychiatry and clinical neurosciences 74, 91–98.

Yizhar, O., Fenno, L.E., Prigge, M., Schneider, F., Davidson, T.J., O’Shea, D.J., Sohal, V.S., Goshen, I., Finkelstein, J., Paz, J.T., Stehfest, K., Fudim, R., Ramakrishnan, C., Huguenard, J.R., Hegemann, P., Deisseroth, K., 2011. Neocortical excitation/inhibition balance in information processing and social dysfunction. Nature 477, 171–178.

Young, K.S., Parsons, C.E., Stein, A., Kringelbach, M.L., 2015. Motion and emotion: depression reduces psychomotor performance and alters affective movements in caregiving interactions. Frontiers in behavioral neuroscience 9, 26.

Yu, P., An, S., Tai, F., Zhang, X., He, F., Wang, J., An, X., Wu, R., 2012. The effects of neonatal paternal deprivation on pair bonding, NAcc dopamine receptor mRNA expression and serum corticosterone in mandarin voles. Hormones and behavior 61, 669–677.

Yuan, Y., Wu, W., Chen, M., Cai, F., Fan, C., Shen, W., Sun, W., Hu, J., 2019. Reward Inhibits Paraventricular CRH Neurons to Relieve Stress. Current biology : CB 29, 1243–1251 e1244.

Zhao, S., Ting, J.T., Atallah, H.E., Qiu, L., Tan, J., Gloss, B., Augustine, G.J., Deisseroth, K., Luo, M., Graybiel, A.M., Feng, G., 2011. Cell type-specific channelrhodopsin-2 transgenic mice for optogenetic dissection of neural circuitry function. Nature methods 8, 745–752.

